# Single cell RNA-seq uncovers the nuclear decoy lincRNA PIRAT as a regulator of systemic monocyte immunity during COVID-19

**DOI:** 10.1101/2021.11.05.467458

**Authors:** Marina Aznaourova, Nils Schmerer, Harshavardhan Janga, Zhenhua Zhang, Kim Pauck, Judith Hoppe, Sarah M Volkers, Daniel Wendisch, Philipp Georg, Margrit Gündisch, Elisabeth Mack, Chrysanthi Skevaki, Christian Keller, Christian Bauer, Wilhelm Bertrams, Andrea Nist, Thorsten Stiewe, Achim D Gruber, Clemens Ruppert, Yang Li, Holger Garn, Leif E Sander, Bernd Schmeck, Leon N Schulte

**Author notes:** These authors contributed equally to this work (Marina Aznaourova, Nils Schmerer).

## Abstract

The systemic immune response to viral infection is shaped by master transcription factors such as NFκB or PU.1. Although long non-coding RNAs (lncRNAs) have been suggested as important regulators of transcription factor activity, their contributions to the systemic immunopathologies observed during SARS-CoV-2 infection have remained unknown. Here, we employed a targeted single-cell RNA-seq approach to reveal lncRNAs differentially expressed in blood leukocytes during severe COVID-19. Our results uncover the lncRNA PIRAT as a major PU.1 feedback-regulator in monocytes, governing the production of the alarmins S100A8/A9 – key drivers of COVID-19 pathogenesis. Knockout and transgene expression, combined with chromatin-occupancy profiling characterized PIRAT as a nuclear decoy RNA, diverting the PU.1 transcription factor from alarmin promoters to dead-end pseudogenes in naïve monocytes. NFκB-dependent PIRAT down-regulation during COVID-19 consequently releases a transcriptional brake, fueling alarmin production. Our results suggest a major role of nuclear noncoding RNA circuits in systemic antiviral responses to SARS-CoV-2 in humans.

## Introduction

Severe courses of infection often culminate in deregulated host responses, ranging from overproduction of inflammation mediators to immune-paralysis (1, 2). Hyperinflammatory and exhausted immune cell states frequently coexist, which poses a challenge to therapeutic interventions. During infections with the pandemic coronavirus SARS-CoV-2, elevated serum levels of NFκB-dependent pro-inflammatory interleukins repeatedly coincide with deranged type I interferon immunity and signs of immune-exhaustion (3). Single-cell RNA-seq (scRNA-seq) studies of peripheral blood mononuclear cells (PBMCs) from patients with severe coronavirus disease 2019 (COVID-19) additionally uncovered a dysregulated myeloid leukocyte compartment, comprising monocytes and granulocytes, and increased production of the PU.1 transcription factor dependent alarmins S100A8 and S100A9 (4-6). The nuclear circuits driving these complex immune rearrangements in COVID-19 remain poorly understood.

To counteract misguided myeloid leukocyte responses, mammalian immune systems have evolved sophisticated control mechanisms, keeping immune gene expression within tight limits at all stages of protein biosynthesis and function. Examples are immune-modulatory splice-regulators, such as the SF3B snRNP (7, 8) or regulators of signaling complex assembly and turnover, such as Optineurin (9). Besides proteins, long non-coding RNAs (lncRNAs) are increasingly recognized as important regulators of mammalian immune responses. Defined as non-coding transcripts ≥ 200 nts, lncRNAs constitute a highly heterogeneous category of RNA, participating in protein complex assembly, disintegration and turnover in the cytoplasmic and nuclear compartment (10-12). So far, only a minor fraction of the ∼20.000 annotated human lncRNAs has been characterized mechanistically and their roles in the human immune system are only beginning to be explored (12). Among the few characterized lncRNAs in this context is MaIL1, which associates with the ubiquitin-reader OPTN to promote TBK1-kinase dependent IRF3 phosphorylation and thus type I interferon immunity (12). GAPLINC, PACER and CARLR regulate pro-inflammatory gene expression by adjusting NFκB p50/p65 expression and activity (13-15). Despite the emerging roles of the noncoding RNA layer in mammalian immunity however (16), the exploration of lncRNA mechanisms contributing to the devastating immune-imbalances during SARS-CoV-2 infection and severe COVID-19 has lagged behind.

Here, we used single cell RNA-seq to study lncRNAs involved in the systemic immunopathologies during COVID-19. Our results highlight the novel lncRNA PIRAT as major regulator of exacerbated PU.1-dependnet alarmin production during SARS-CoV-2 infection. A single nucleotide polymorphism in the PIRAT locus has recently been associated with hematological malignancies (17), the function of PIRAT, however had remained unknown. Using CRISPR/Cas9, lentiviral gene transfer and global chromatin occupancy profiling, we show that PIRAT functions to redirect the PU.1 transcription factor from alarmin promoters to dead-end pseudogenes. This serves to suppress PU.1-dependent S100A8 and S100A9 alarmin expression under homeostatic conditions. Down-regulation of PIRAT in patients with severe COVID-19 consequently removes a transcriptional break on alarmin production. These results uncover a hitherto unknown RNA-layer, governing peripheral monocyte activation and PU.1-dependent systemic alarmin production – hallmarks of severe COVID-19.

## Results

### Identification of COVID-19 relevant myeloid lincRNA signatures

To chart candidate long intergenic noncoding RNAs (lincRNAs) relevant to the disturbed myeloid immune compartment in COVID-19, we consolidated RNA-seq data from several sources, followed by in-depth scRNA-seq profiling (Fig. 1A). At first, leukocyte-specific mRNAs and lincRNAs were narrowed down using Illumina Human Bodymap data (Fig. 1A). Confirming successful extraction of leukocyte-specific RNAs from these datasets, pathway analysis revealed an exclusive enrichment of immune-relevant terms (Fig. S1A and B). We then charted expression of these transcripts amongst three publicly available replicates of peripheral blood monocyte, granulocyte, B-cell, NK-cell and T-cell RNA-seq profiles (18, 19). PCA and hierarchical clustering analysis successfully discriminated the major leukocyte compartments, based on their lincRNA and mRNA profiles, respectively (Fig. 1B, Fig. S1C and D). To confirm the cell-type specificity of the interrogated myeloid and lymphoid lincRNAs, we studied their expression in purified peripheral blood derived macrophages, dendritic cells, monocytes, granulocytes, NK cells, B cells and naïve (CD45RO^-^) or memory (CD45RO^+^) T-cells. qRT-PCR confirmed the preferential expression of LINC00211 (henceforth PIRAT, for PU.1-induced regulator of alarmin transcription), LUCAT1 and AC064805.1 in myeloid cells, whereas LINC02295, LINC02446 and LINC00861 were confirmed as lymphoid cell specific transcripts (Fig. 1C and Fig. S1E-I). Of note, among the lymphoid lincRNAs, LINC02446 was particularly abundant in CD8^+^/CD45RO^+^ T cells, indicating a specific role in the CD8-memory T-cell niche (Fig. S1H and I). Among the myeloid lincRNAs, our attention was caught by PIRAT, since a SNP in the PIRAT locus (rs4670221-G, p-value: 3 × 10^−10^) had recently been associated with haematological alterations (17). The function of PIRAT, however, had remained unknown. Besides PIRAT, LUCAT1 was selected as a candidate lncRNA relevant to myeloid immunity in COVID-19 due to its particularly high expression in monocytes and granulocytes.

**Fig. 1.**
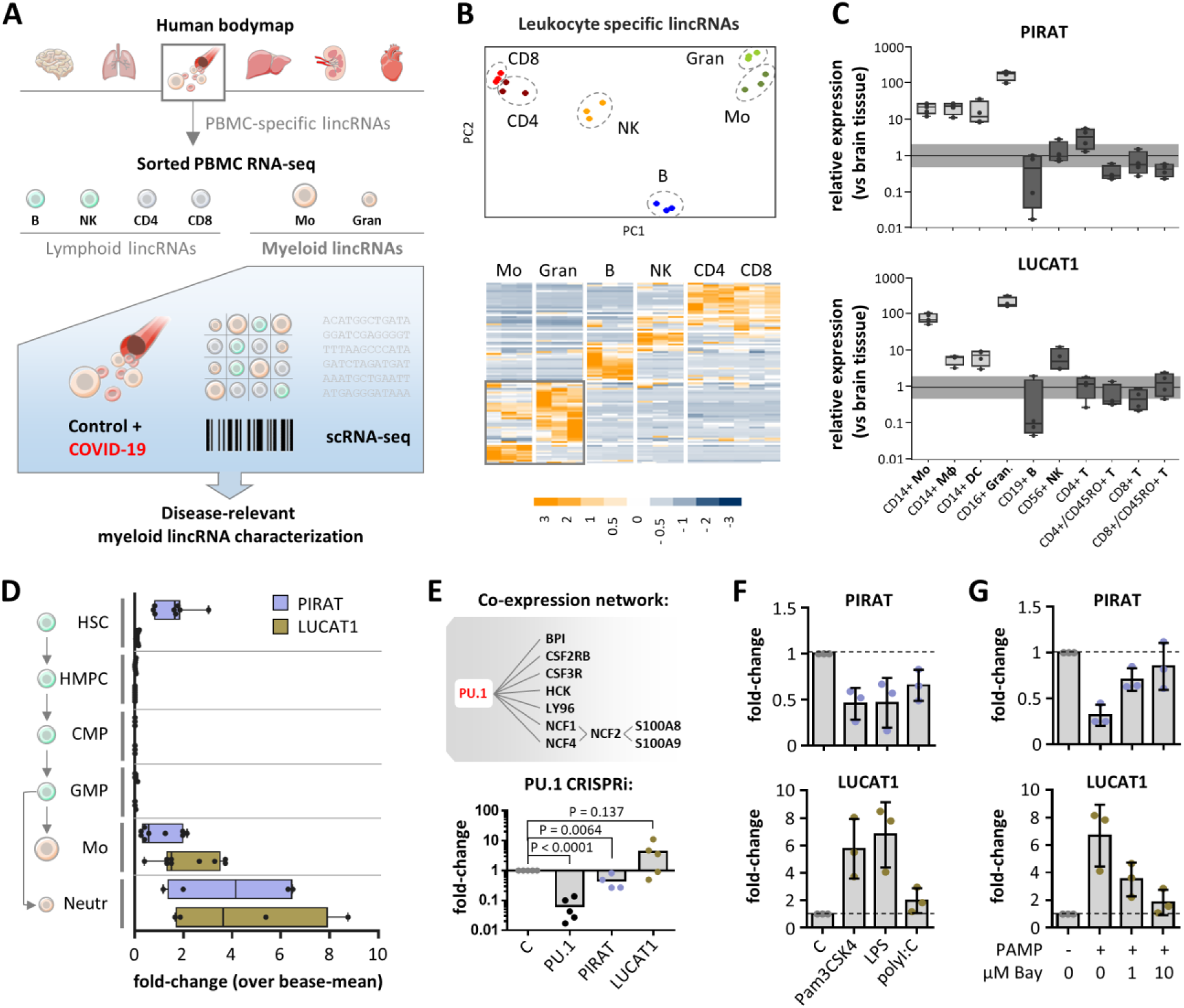
Identification of human myeloid lineage-specific lincRNAs. A) Bulk and scRNA-seq analysis strategy for the determination of myeloid lincRNAs relevant to COVID-19. B) PCA (top) and hierarchical clustering (bottom, row Z-scores) of monocyte, granulocyte, B-, NK- and CD4^+^ and CD8^+^ T-cell lincRNA expression profiles. C) qRT-PCR validation of PIRAT (LINC00211) and LUCAT1 as myeloid-specific lincRNAs (expression relative to human brain tissue). Horizontal bar indicates base-line (black) and 2-fold deviation from base-line (grey). D) Relative abundance of PIRAT and LUCAT1 in RNA-seq profiles of hematopoietic stem cells (HSC), hematopoietic multipotent precursor cells (HMPC), common myeloid progenitors (CMP), granulocyte-macrophage progenitors (GMP), monocytes and neutrophils. E) Top: PIRAT co-expression network; bottom: qRT-PCR analysis of PU.1 mRNA, PIRAT and LUCAT1 expression in PU.1 knockdown compared to control THP1 monocytes. F-G) qRT-PCR analysis of lincRNA expression in response to indicated PAMPs and NFκB inhibitor BAY-11-7082 (PAMP = 4 h LPS + polyI:C stimulation). C-G: ≥ 3 independent experiments and one-way ANOVA test.

To determine at which stages of myeloid ontogeny both lincRNAs become relevant, we traced their expression from haematopoietic stem- and progenitor-cells to mature leukocytes, using Blueprint Consortium RNA-seq profiles (20). Expression of PIRAT declined upon haematopoietic stem cell (HSC) differentiation into multipotent progenitors (HMPCs) and, similar to LUCAT1, remained low during the common myeloid progenitor (GMP) and granulocyte-monocyte progenitor (GMP) stages (Fig. 1D). Expression of both lincRNAs, however, strongly increased in mature monocytes and neutrophils (Fig. 1D). Interestingly, co-expression analysis suggested PIRAT to depend on an expression network driven by the myeloid master transcription factor PU.1 (Fig. 1E, Fig. S2). Among the PIRAT-co-expressed genes were the PU.1-dependent major alarmins S100A8 and S100A9 (Fig. 1E, Fig. S2), which play a key role in COVID-19 pathogenesis (4-6, 21, 22). Dependence of PIRAT but not LUCAT1 on the PU.1 transcription factor was confirmed by PU.1 knock-down in THP1 monocytes (Fig. 1E). Further underscoring their differential dependence on myeloid expression programs, PIRAT was down- and LUCAT1 was up-regulated in an NFκB-dependent manner upon monocyte activation with immune agonists (Fig. 1F and G). Thus, PIRAT and LUCAT1 are myeloid signature lncRNAs, activated during late haematopoiesis and differentially depending on the immune master-regulators PU.1 and NFκB.

### Single-cell resolved myeloid lincRNA responses to SARS-CoV-2 infection

Recent scRNA-seq studies have revealed profound changes in myeloid coding gene expression networks in severe COVID-19. To dissect the contributions of myeloid signature lncRNAs, such as PIRAT and LUCAT1 to these alterations, we performed BD Rhapsody scRNA-seq analysis of PBMCs from control and severe COVID-19 patients (WHO grade ≥ 5) using an immune-response panel combined with a custom lncRNA detection panel (Fig. 2A). qRT-PCR confirmed the expected induction of the immune-response markers CXCL2 and IL6 in COVID-19 patients (Fig. 2B). ScRNA-seq analysis of patient PBMCs charted all expected myeloid and lymphoid cell populations and discriminated 4 monocyte populations along the CD14-, CD16- and HLA-expression scheme (Fig. 2C and D, Fig. S3A-D). FACS analysis confirmed the previously reported increased abundance of immature CD15^++^/CD24^++^ neutrophils in peripheral blood from patients with severe COVID-19, as well as the reduction of CD14^++^/CD16^dim^ classical monocytes during mild and severe COVID-19, indicative of myeloid exhaustion (4) (Fig. 2E). Differential gene expression and Reactome pathway analysis confirmed the pro-inflammatory activation of classical, non-classical and intermediate monocytes during COVID-19 (Fig. 2F and Fig. S4A-D). Analysis of lincRNA distribution in the scRNA-seq profiles confirmed the expected abundance of the B-cell proliferation promoting lncRNA BIC (23) in B-lymphocytes. Furthermore, BIC was up-regulated in dendritic cells from COVID-19 compared to control patients, in line with its known role in antigen presenting cell (APC) activation (Fig. 2G) (24). Moreover, we observed the expected induction of the type I interferon inducing lncRNA MaIL1 (12) in all monocyte populations, but also in B-cells from SARS-CoV-2 infected patients (Fig. 2G). Intriguingly, the scRNA-seq data also confirmed the strict myeloid expression pattern of LUCAT1 and PIRAT and revealed their preferential expression in CD14^+^-monocytes. Unlike in classical and intermediate monocytes, expression remained low in non-classical CD16^+^-monocytes (Fig. 2G). Whereas LUCAT1 expression was up-regulated in classical and intermediate monocytes during COVID-19, PIRAT was downregulated, reminiscent of the differential regulation of both lincRNAs in response to sterile immune agonists (Fig. 2G compared to Fig. 1F-G). Preferential expression of both lincRNAs in classical rather than non-classical monocytes and their opposite regulation during COVID-19 was confirmed in cell sorting and qRT-PCR validation experiments (Fig. 2H and I). Taken together, these results confirm an imbalanced myeloid leukocyte compartment during severe COVID-19 and reveal LUCAT1 and PIRAT as CD14^+^-monocyte-specific myeloid lincRNAs, up- and down-regulated during SARS-CoV-2 infection, respectively.

**Fig. 2.**
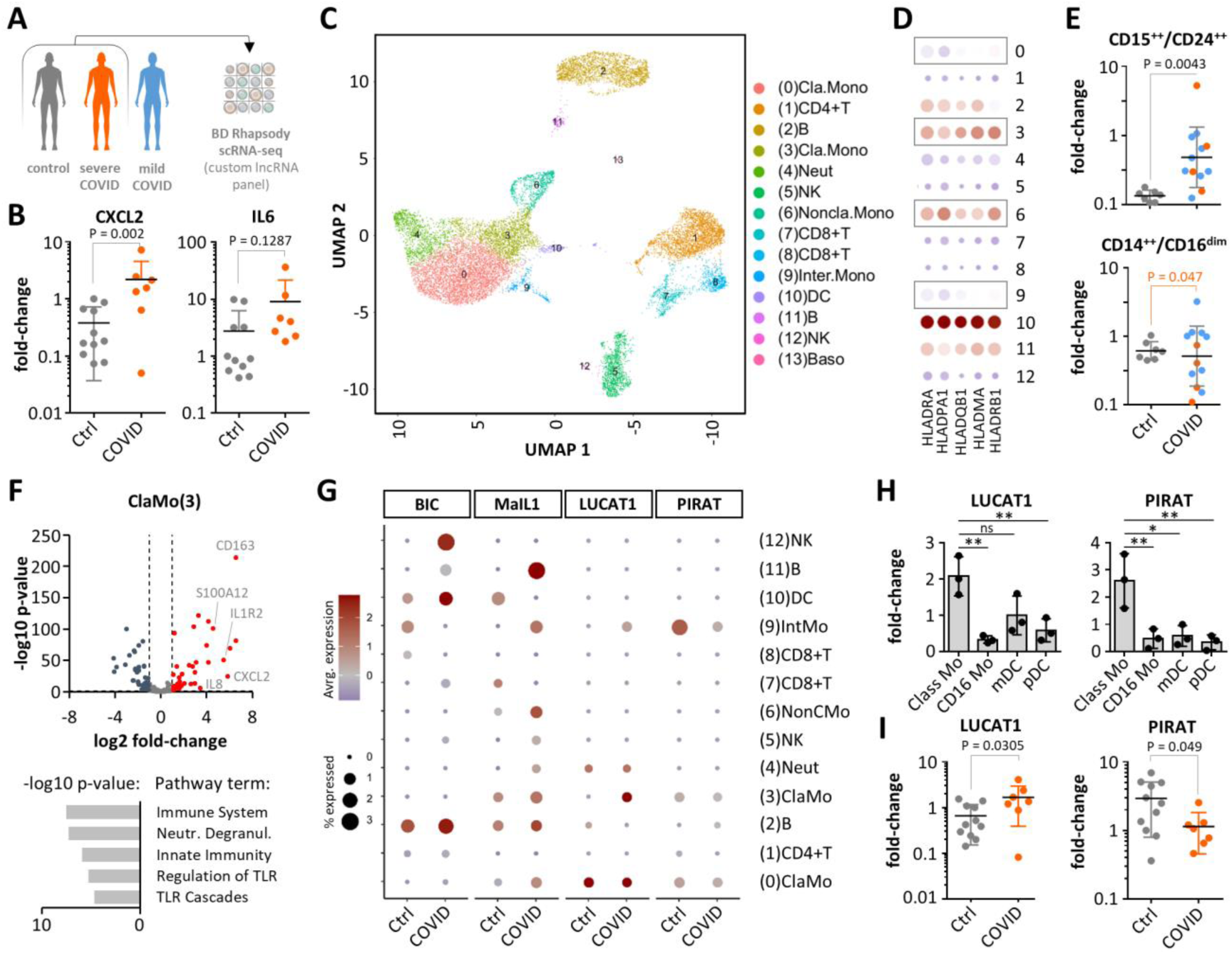
Single cell RNA-seq analysis of lincRNA expression during COVID-19. A) Control and COVID-19 patient PBMC analysis strategy. B) Validation of immune marker induction in COVID-19 cohort whole PBMCs (qRT-PCR, control-patient 1 set as reference). C) UMAP-plot with color-coded cell populations identified in merged scRNA-seq data. D) HLA mRNA expression profile (scRNA-seq, monocytes highlighted). E) FACS validation of immature neutrophil (CD15^++^/CD24^++^) appearance and reduction of classical monocytes (CD14^++^/CD16^dim^) in COVID-19 patient blood (color-coded according to A). F) Volcano plot (top) and Reactome pathway (bottom) analysis of classical monocyte response during COVID-19 (scRNA-seq). G) Cell-type specific lincRNA profiles in control and COVID-19 patients (scRNA-seq). H) PIRAT and LUCAT1 expression in classical (Class) and non-classical (CD16) monocytes, myeloid dendritic cells (mDC) and plasmacytoid DCs (pDC) (qRT-PCR). I) Same as B, but for PIRAT and LUCAT1. B, E, I: Student’s t-test. H: One-way ANOVA, 3 independent experiments.

### COVID-suppressed lincRNA PIRAT antagonizes alarmin expression in monocytes

While our manuscript was in preparation, LUCAT1 was reported to act as a negative feedback regulator of interferon-responses in human myeloid cells (25). Thus, the up-regulation of LUCAT1 in monocytes during severe COVID-19 (Fig. 2G and I) likely reflects the activation of systemic interferon immunity. Since the function of PIRAT had remained unknown, we decided to focus on the role of this lincRNA in human monocytes in the present study and to decipher the reasons for its down-regulation during COVID-19. As current lncRNA annotations have to be regarded as provisional, we mapped the exact PIRAT transcript architecture by RACE-PCR. Deviating from the current GENCODE annotation, 5’ and 3’ RACE revealed a 2-exon structure in primary human monocytes (Fig. 3A, Fig. S5). ENCODE monocyte RNA-seq, DNaseI-seq and ChIP-seq data confirmed a DNaseI hyper-sensitive site at the mapped PIRAT 5’-end and H3K4 trimethylation and RNA-seq coverage across the RACE-refined gene body – hallmarks of transcriptionally active regions (Fig. 3A). Re-analysis of the refined PIRAT sequence by the CPC2 algorithm confirmed low protein-coding potential, similar to the well-established non-coding RNAs XIST and HOTAIR and different from mRNAs (ACTB, GAPDH, IL1B) (Fig. 3B). PIRAT cDNA sequence conservation exceeded 90 % in the genomes of catarrhine primates but dropped to 33.5 % in mice (Fig. 3C). Thus, PIRAT is a 2-exon lincRNA, stably maintained during higher primate evolution.

**Fig. 3.**
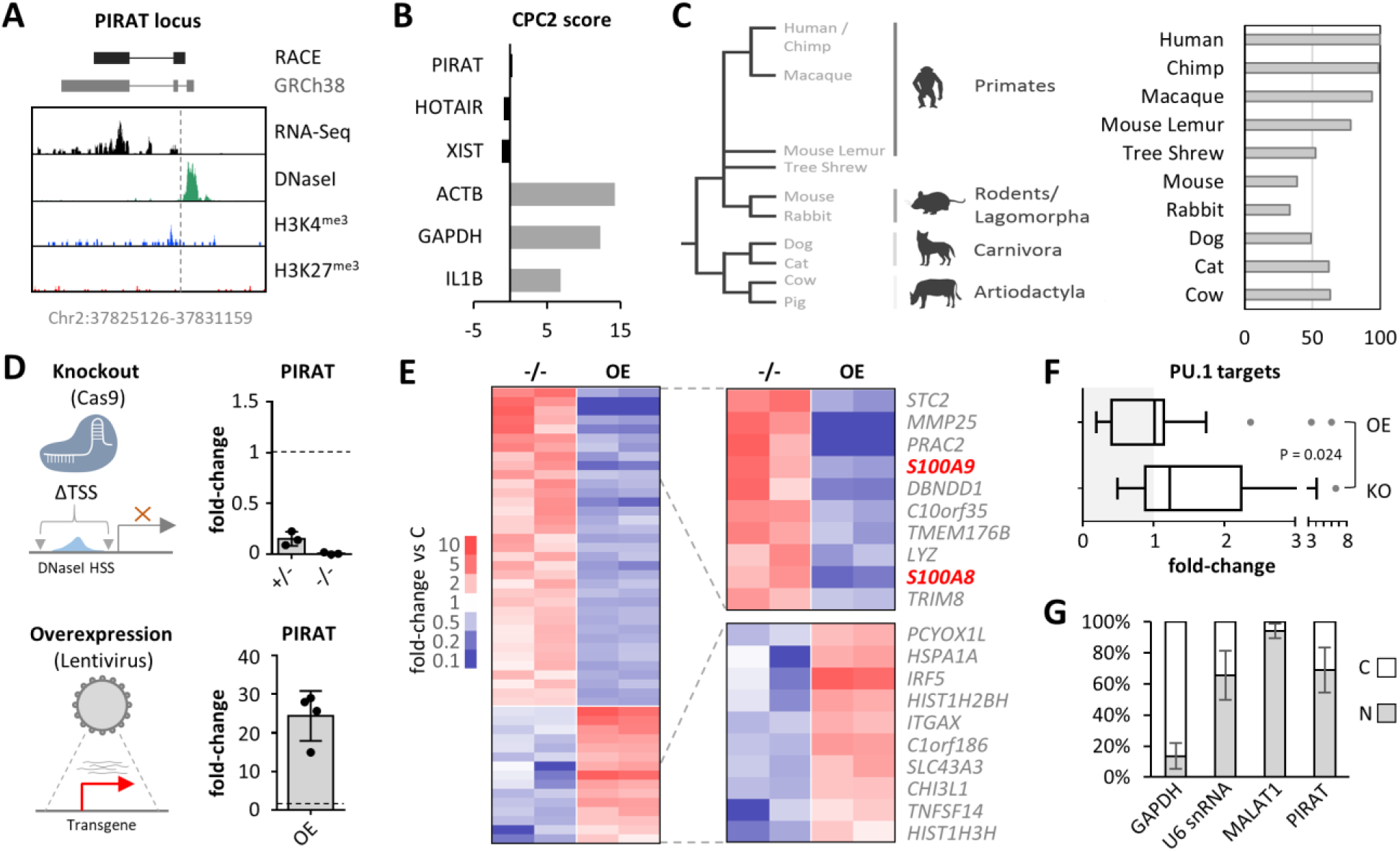
Role of nuclear lincRNA PIRAT in human monocytes. A) RACE-PCR refined (black) and annotated (grey) PIRAT splice structure and chromosomal position, compared to matched ENCODE human primary CD14^+^-monocyte RNA-Seq, DNaseI-Seq, and ChIP-Seq (H3K4me3 and H3K27me3) coverage. B) CPC2 coding score of indicated lncRNAs and mRNAs. For PIRAT, the RACE-PCR refined cDNA sequence from A) was used. C) Left: phylogenetic relationship of selected mammalian species and their respective orders. Right: conservation of the RACE-PCR refined PIRAT cDNA sequence in the genomes of the respective species. D) Representation and qRT-PCR-validation of PIRAT mono- (+/-) and biallelic (-/-) knockout and lentiviral over-expression strategy (THP1 monocytes). E) RNA-seq analysis of PIRAT knockout (-/-) and overexpression (OE) cells (color-coded mRNA fold-changes ≥ 2, compared to wild-type cells). F) Base-mean fold-changes of PU.1 target genes in datasets from E. G) Subcellular localization of PIRAT in primary CD14^+^-monocytes (qRT-PCR, three independent experiments; C = cytoplasm, N = nucleus). F: Student’s t-test.

To reveal the cellular function of the lincRNA, we generated PIRAT promoter-deficient THP1 monocytes using CRISPR/Cas9 (Fig. 3D, Fig. S6A). Additionally, we generated cells overexpressing PIRAT from a lentiviral backbone (Fig. 3D). Global expression profiling by poly(A)-RNA-seq uncovered dozens of mRNAs regulated (≥ 2-fold) into opposite directions upon PIRAT knockout and overexpression, respectively (Fig. 3E). Intriguingly, among the top 10 PIRAT-suppressed genes were the PU.1-dependent alarmins S100A8 and S100A9, which are also co-expressed with PIRAT in myeloid cells (Fig. 3E compared to Fig.1E) and promote COVID-19 pathogenesis (4, 5, 21, 22). Beyond S100A8 and A9, the suppressive effect of PIRAT extended to other direct PU.1 target genes (Fig. 3F). Reciprocally, genes suppressed by PU.1, such as ITGAX (CD11c) or CHI3L1 (26-28), were de-repressed upon PIRAT knockout (Fig. 3E). Subcellular fractionation qRT-PCR characterized PIRAT as a nuclear-retained lincRNA (Fig. 3G), which was further corroborated by RNA-FISH (Fig. S6B). Thus, PIRAT is a myeloid nuclear RNA, down-regulated in CD14^+^-monocytes during COVID-19 to promote PU.1-dependent alarmin expression.

To gain further insights into the physiological role of PIRAT during COVID-19, we overlaid the RNA-seq profiles of PIRAT-manipulated cell lines with the scRNA-seq profiles of COVID-19 and control patient PBMCs. Among all mRNAs up- or down-regulated ≥ 2-fold during COVID-19 (scRNA-seq data, classical monocytes) or upon PIRAT expression-manipulation (THP1 monocytes), 33 were detected in both datasets. The overlap of mRNAs regulated ≥ 2-fold in both datasets was 12.1 % (4 mRNAs, Fig. 4A). Among the mRNAs up-regulated during COVID-19 (scRNA-seq data), S100A8 and S100A9 experienced the strongest de-repression upon PIRAT knockout (Fig. 4B and C). Vice versa, genes down-regulated in CD14^+^-monocytes during COVID-19 were under significant positive influence by PIRAT, headed by the PU.1-suppressed genes IRF5 and ITGAX (Fig. 4B and C). Thus, genes activated and suppressed by PIRAT are reciprocally regulated by PU.1 and in COVID-19. This notion was further corroborated in qRT-PCR and FACS validation experiments, which confirmed the control of S100A8, S100A9 and ITGAX expression by PIRAT and the regulation of these factors during COVID-19 (Fig. 4D, E, F, G and Fig. S6C, D). Finally, knockdown of PU.1 in THP1 monocytes using CRISPR interference, verified the dependence not only of PIRAT but also of the alarmins S100A8 and S100A9 on this transcription factor (Fig. 4H, Fig. 1E). Taken together, this suggests PIRAT to function as a negative feedback regulator of PU.1, limiting S100A8 and A9 alarmin expression in monocytes at base-line. NFκB-dependent down-regulation of PIRAT (Fig. 1G) consequently removes a molecular break on the production of alarmins known to contribute to the development of severe COVID-19.

**Fig. 4.**
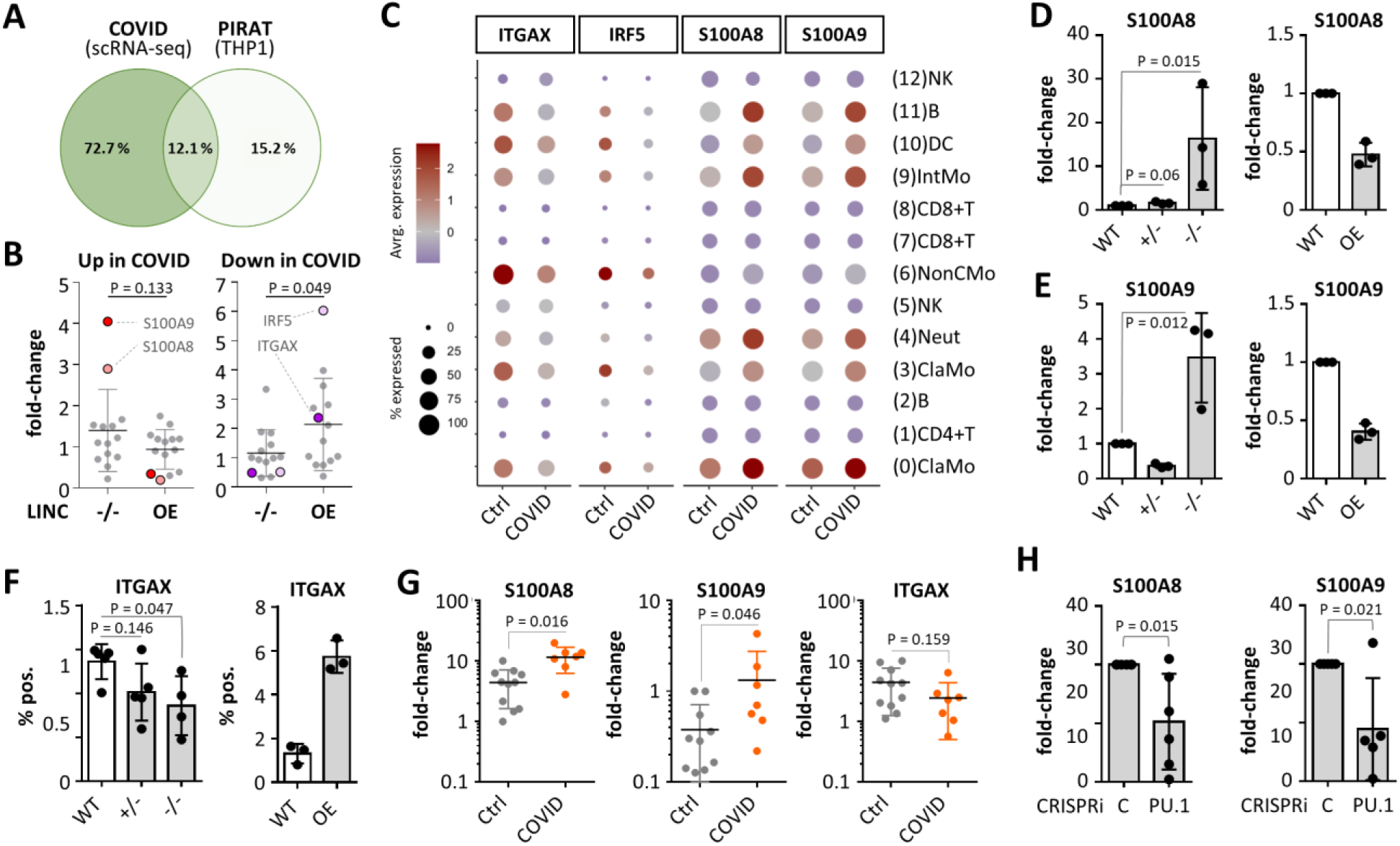
Participation of PIRAT in monocyte PU.1 circuits relevant to COVID-19. A) Overlap of genes regulated ≥ 2-fold in classical monocytes during COVID-19 (scRNA-seq data) and upon PIRAT knockout / over-expression (data from Fig. 3E). B) Regulation of genes in PIRAT knockout (-/-) or over-expression (OE) compared to wild-type cells, up-regulated (left) or down-regulated (right) during COVID-19. C) Cell-type specific expression of top PIRAT-controlled, COVID-responsive mRNAs from B (scRNA-seq). D-F): Expression changes of PIRAT-controlled PU.1 target genes in PIRAT knockout and overexpression compared to wild-type THP1 monocytes (qRT-PCR validations). G) Regulation of PIRAT-controlled genes in PBMC samples from COVID-19 and control-patients (qRT-PCR, control-patient 1 set as reference). H) Regulation of S100A8 and S100A9 in PU.1 knockdown compared to control THP1 monocytes. D-F: One-way ANOVA; G-H: Student’s t-test, ≥ 3 independent experiments.

### PIRAT redirects PU.1 from alarmin gene promoters to pseudogenes

To interrogate the molecular mechanism of alarmin control by PIRAT in the nucleus, we investigated the physical interaction of this lincRNA with chromatin and PU.1 using RNA and protein affinity chromatography. Antisense-purification of PIRAT-occupied chromatin from primary CD14^+^-monocyte lysates by ChIRP (Fig. 5A) recovered PIRAT RNA and verified the expected crosslinking of PIRAT to its own genomic site of transcription (Fig. 5B and C). Refusing a model where the lincRNA controls the PU.1 transcription factor directly at its target gene promoters, PIRAT did not bind to the PU.1 occupied region upstream to the S100A8 alarmin gene (Fig. 5D, Fig. S7A). In search of alternative mechanistic explanations for the observed negative feedback control of PU.1 by PIRAT, we recorded the global genome occupancy profile of PIRAT in CD14^+^-monocytes using ChIRP-seq. Unexpectedly, peak-calling revealed PIRAT to occupy multiple sites along the uncharacterized REXO1L-pseudogene array at chromosome 8q21.2 (Fig. 5E). Importantly, comparison of the PIRAT ChIRP-seq profile with matched ENCODE CD14^+^-monocyte ChIP-seq data uncovered a repetitive binding pattern composed of alternating PIRAT and PU.1 binding sites along the entire open chromatin of the REXO1LP pseudogene repeat (Fig. 5F). PIRAT and PU.1 binding to selected peaks within the REXO1LP array was confirmed by PIRAT ChIRP-qRT-PCR and PU.1 ChIP-qRT-PCR (Fig. 5G and H). We therefore hypothesized that PIRAT buffers PU.1 activity by tethering the transcription factor to multiple binding sites within the REXO1L-pseudogene repeat (Fig. 5I). In line with this model, we observed a direct interaction of PIRAT with PU.1 in primary monocytes in UV-CLIP experiments (∼ 12-fold enrichment, Fig. 6A). ChIP confirmed the binding of PU.1 to the promoters of S100A8 and A9 in primary monocytes (Fig. 6B, Fig. S7A) and PU.1-binding to these promoters was enhanced in PIRAT-deficient compared to wild-type THP1 monocytes (Fig. 6C). Concurrently, PU.1-binding to the REXO1L-pseudogene repeat was diluted in the absence of PIRAT (Fig. 6C). We therefore propose a function of the REXO1L-pseudogene array as a PIRAT-dependent cellular cache for PU.1. PIRAT downregulation in classical CD14^+^-monocytes under infectious conditions renders this cached PU.1 pool accessible and propels alarmin production during severe COVID-19 (Fig. 6D).

**Fig. 5.**
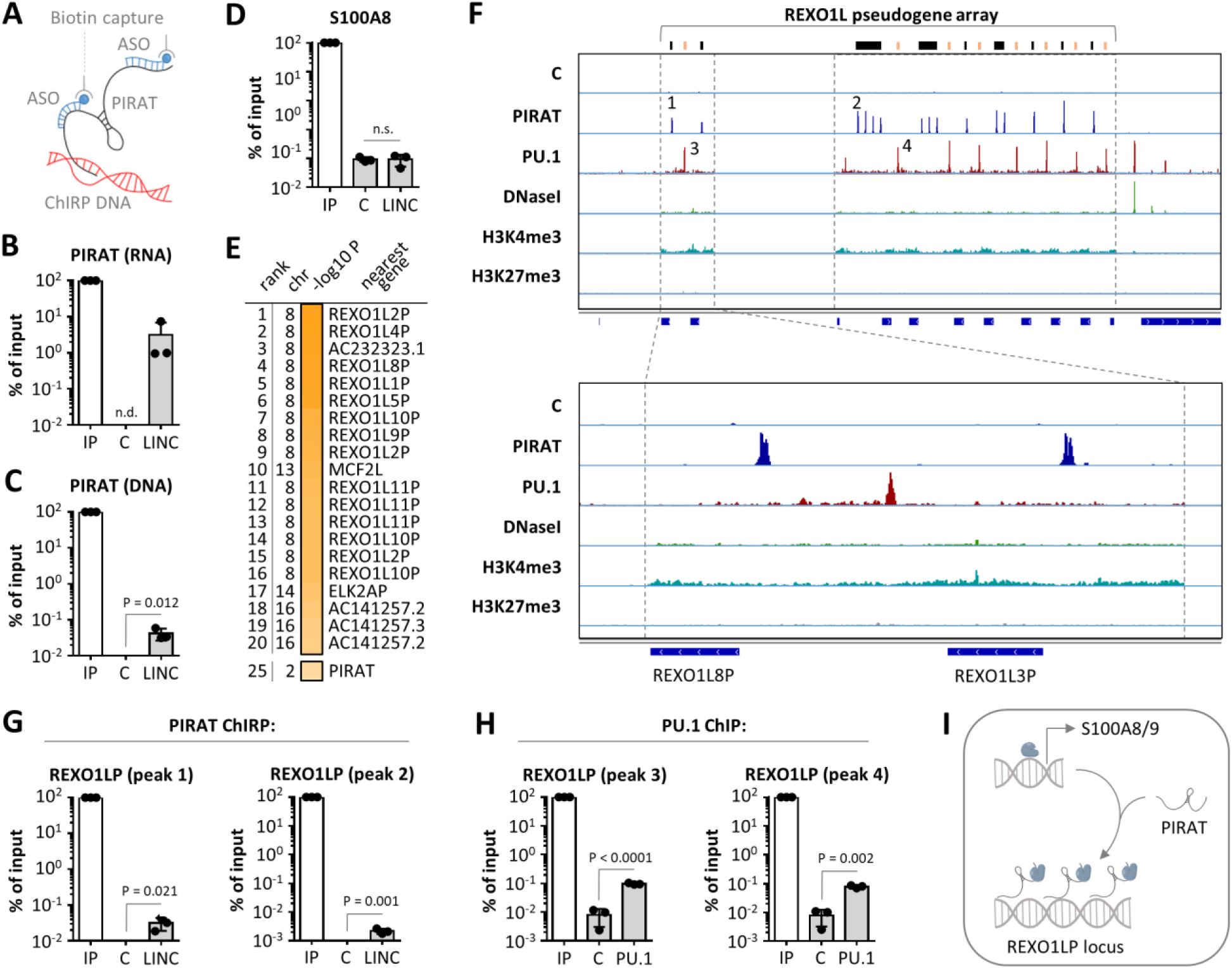
Repetitive binding of PU.1 and PIRAT to the REXO1LP pseudogene locus. A) PIRAT ChIRP was performed as illustrated, using primary human CD14^+^-monocytes. B) Recovery of PIRAT RNA in ChIRP samples compared to input (IP) sample (C = control ChIRP, LINC = PIRAT ChIRP; qRT-PCR analysis). C) Same as B but with genomic DNA. D) Same as C but with S100A8 promoter detection. E) Summary of PIRAT binding site peak-calling (ChIRP-seq; chr = chromosome; top 20 peaks and peak # 25 are shown). F) IGV plots showing control (C) and PIRAT ChIRP-seq and matched CD14^+^-monocyte PU.1 ChIP-, DNaseI- and histone-3 ChIP-seq coverage in the REXO1L pseudogene locus. G) qRT-PCR validation of PIRAT binding to ChIRP-seq peaks indicated in F. H) qRT-PCR validation of PU.1 binding to ChIP-seq peaks indicated in F. I) Model of PU.1 redirection from alarmin promoters to the REXO1LP locus by PIRAT. B,C,D,G,H: Student’s t-test, three independent experiments.

**Fig. 6.**
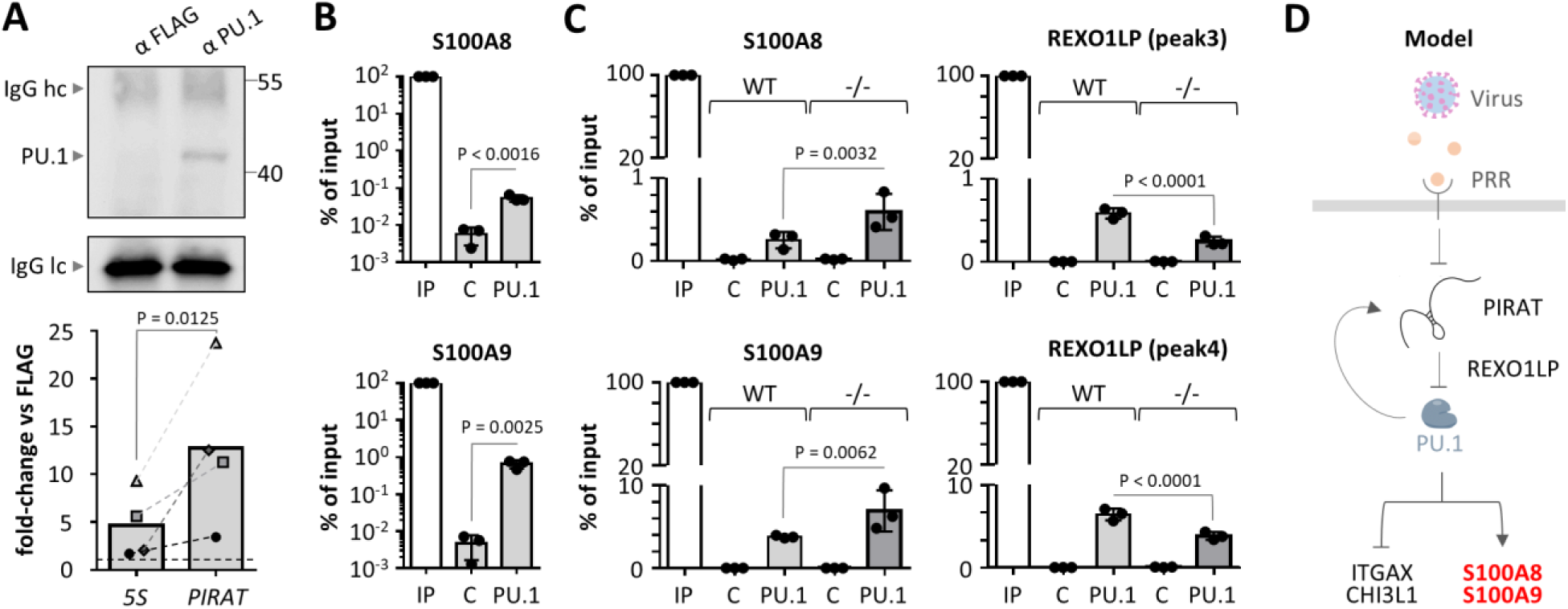
PIRAT redirects PU.1 from alarmin promoters to the REXO1LP repeat. A) Western blot validation of PU.1 capture (top; hc/lc = light chain/heavy chain) and qRT-PCR analysis of PIRAT enrichment (bottom) in PU.1 UV-CLIP-experiments (primary CD14^+^-monocytes). B) ChIP qRT-PCR analysis of PU.1 binding to S100A8 and A9 promoter DNA in primary CD14^+^-monocytes. C) qRT-PCR analysis of PU.1 binding to ChIP-seq peaks in the S100A8 and S1009 promoters and in the REXO1LP pseudogene locus, in wild-type (WT) and PIRAT knockout (-/-) THP1 monocytes. D) Summary of negative feedback function of PIRAT during PU.1-dependent alarmin control and infection. A-B: Student’s t-test. C: One-way ANOVA. ≥ 3 independent experiments were performed.

### PIRAT is a myeloid marker with clinical utility beyond COVID-19

Given the highly specific expression and function of PIRAT in myeloid immune cells, we predicted its utility as a general marker of myeloid cell abundance and tissue infiltration in COVID-19 and other infectious and inflammatory diseases. In line with this hypothesis, the expression of PIRAT in PBMC samples from control and COVID-19 patients significantly correlated with the relative abundance of CD24^+^ neutrophils and classical monocytes, but not with non-classical monocytes (Fig. 7A). This is in agreement with our scRNA-seq results, showing little expression of PIRAT in CD16-positive non-classical monocytes (Fig. 2). Beyond COVID-19, PIRAT levels significantly correlated with the percentage of infiltrating myeloid cells (granulocytes; R^2^ = 0.82) in bronchoalveolar lavage fluid (BALF) from patients with bronchopulmonary infection (Fig. 7B). To additionally test the utility of PIRAT as a marker of myeloid cell infiltration in non-infectious lung diseases, we monitored the levels of the lincRNA in tissue samples from idiopathic pulmonary fibrosis (IPF) patients. Neutrophils play an important role in IPF tissue remodeling and elevated migration of these cells into IPF tissue has been associated with early mortality (29). Similar to the results with BALF from patients with pulmonary infection, PIRAT levels significantly correlated with the percentage of neutrophils in IPF tissue (R^2^ = 0.83) but not with NK cells (R^2^ = 0.14) (Fig. 7C). Thus, PIRAT is a suitable RNA marker for myeloid immune cell infiltration and abundance in biomaterial from patients with infectious and non-infectious diseases, in line with its important immunoregulatory role in the myeloid system.

**Fig. 7.**
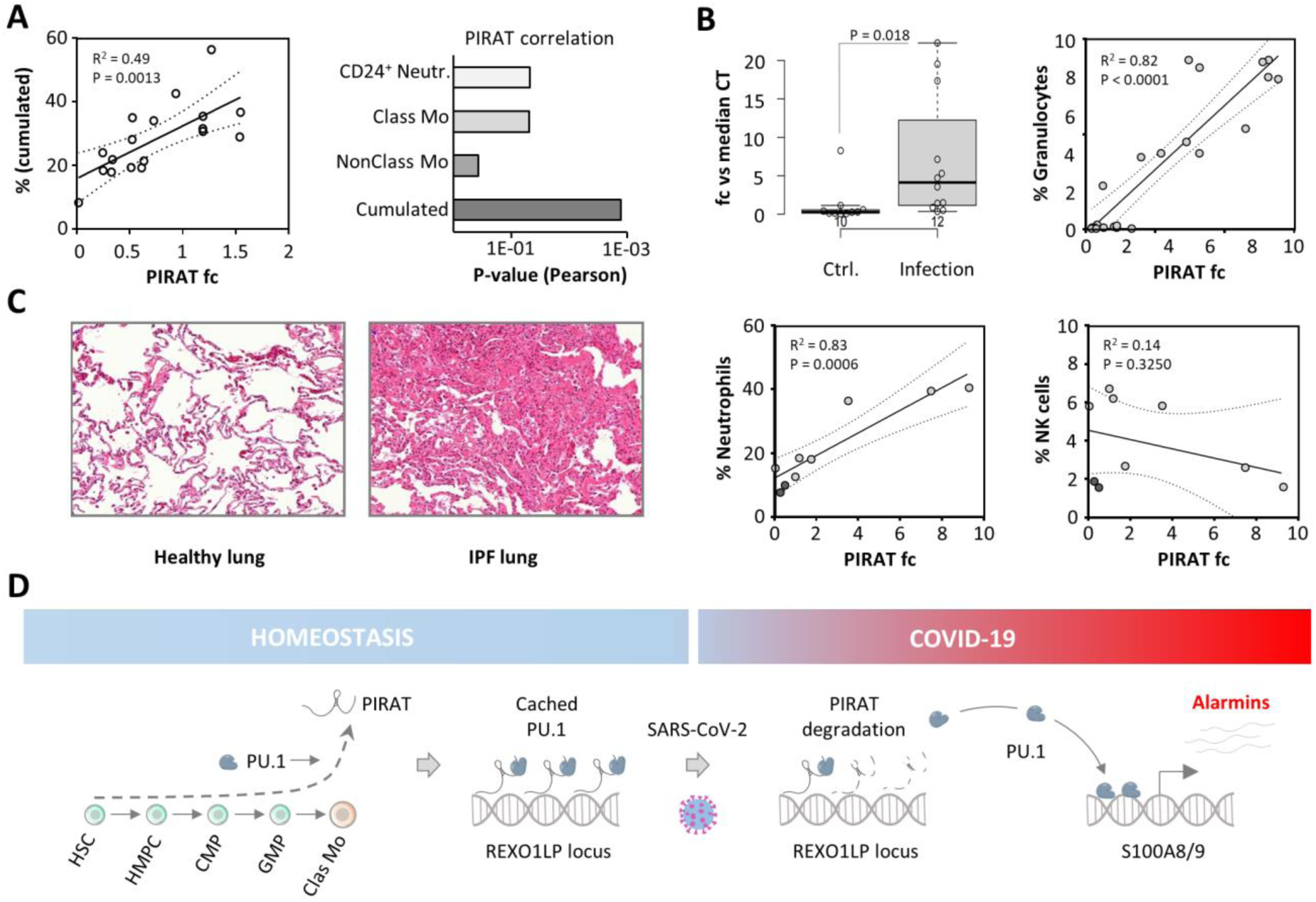
Demarcation of the myeloid compartment by PIRAT in COVID-19 and beyond. A) Left: Pearson correlation of cumulated neutrophil and classical monocyte percentage with PIRAT levels in COVID-19 patient PBMC samples. Right: P-values for PIRAT correlation (Pearson) with the percentages of the indicated cell types in COVID-19 patient PBMC samples. B) Left: qRT-PCR analysis of PIRAT expression in BALF from control and pulmonary infection patients (individual data-points shown; two-tailed Student’s t-test). Right: Pearson correlation of granulocyte percentage with PIRAT expression in BALF samples. C) Left: Representative H&E-stained sections of human healthy and late-stage IPF lung tissue. Right: Pearson correlation of neutrophil or NK cell percentage with PIRAT fold-change (compared to IPF lung # 3). D) Summary of PIRAT function under homeostatic and infection conditions: upon differentiation of bone-marrow precursors into mature myeloid cells, PIRAT expression is activated to tether PU.1 to pseudogenes and restrain PU.1 activity in the myeloid lineage. PIRAT degradation in monocytes during severe COVID- 19 releases inactive PU.1 from the REXO1L pseudogene locus and propels S100A8 and S100A9 alarmin expression. Exaggerated alarmin production contributes to COVID-19 pathogenesis.

In summary, our results suggest a vital contribution of lincRNAs to the systemic immune response to SARS-CoV-2 infection and reveal PIRAT as a novel nuclear RNA, restraining PU.1 dependent S100A8/A9 alarmin production in the myeloid lineage. Activation of PIRAT upon differentiation of myeloid precursors in the bone marrow establishes a break on PU.1 target gene expression by redirecting PU.1 to pseudogenes. During SARS-CoV-2 infection, NFκB-dependent down-regulation of PIRAT releases a break on PU.1 activity and fuels the expression of the major alarmins S100A8 and S100A9, which contribute to myeloid imbalances in COVID-19 (Fig. 7D). We believe that the exploration of further lncRNAs specifically expressed in myeloid and lymphoid immune cells will provide a vastly improved, therapy-relevant understanding of the molecular circuits contributing to the progression of severe inflammatory and infectious diseases.

## Discussion

During SARS-CoV-2 infection, a subset of patients develops a severe course of disease with complex alterations in the peripheral immune system. Besides characteristic cellular changes, indicative of emergency myelopoiesis, severe COVID-19 entails systemic inflammatory components also registered in other difficult to treat infectious disease trajectories (1, 2). A better understanding of the underlying molecular circuits is urgently needed to improve the outcome of infections with SARS-CoV-2 and other potentially pandemic agents. Several recent studies have employed scRNA-seq approaches to dissect peripheral immune alterations in COVID-19 (3-5). So far, however, the noncoding RNA layer has been neglected, despite the emerging role of these RNAs in immune-regulation.

Here, we employed an lncRNA-centric approach to dissect mechanisms underlying immune-alterations in COVID-19 at the single cell level. Our results suggest that the lincRNA PIRAT governs critical myeloid immune-trajectories in COVID-19 at the transcriptional level. PIRAT is primarily expressed in monocytes - a critical source of peripheral immune-mediators, such as S100A8 and A9 in COVID-19 (5). Our finding that PIRAT restrains the production of S100A8 and A9 assigns this lincRNA an important role in myeloid immune responses during severe COVID-19. Upstream of S100A8 and A9, PIRAT controls PU.1 as a negative feedback regulator. Feedback control constitutes a universal regulatory principle, conferring stability to cellular circuits (30). Mechanistically, PIRAT redirects PU.1 from alarmin promoters to the REXO1LP locus, which suggests a novel function of pseudogenes as nuclear caches for transcription factors. Downregulation of PIRAT during COVID-19 releases this cached transcription factor pool. Since PU.1 is a master-regulator of myelopoiesis, PIRAT might also contribute to the imbalanced myeloid differentiation-trajectories seen in severe COVID-19, independent of S100A8 and A9. The PU.1 dose for instance decides over the commitment to the macrophage and granulocyte differentiation paths, respectively (31, 32). Furthermore, reduction of PU.1 levels is required for megakaryocyte differentiation and thus platelet production (33). This might also explain the association of a SNP in the *PIRAT* locus with altered platelet volume (17). Granulocyte and platelet differentiation trajectories again are disturbed in COVID-19 (4, 34). *In vivo* studies could further clarify the role of PIRAT in myeloid cell differentiation and activation. The low sequence conservation of PIRAT in rodents (Fig. 3C), however, calls into question the possibility of such investigations. Beyond PIRAT, further lncRNAs likely regulate the differentiation of myeloid cells. Recently, Schwarzer et al. identified a nuclear lncRNA, *LINC00173*, as a regulator myeloid progenitor proliferation, contributing to granulopoiesis. *LINC00173* negatively impacts the expression of stemness-associated genes, probably through PRC2 complex dependent modifications at HOX-gene loci (35). Similarly, the lncRNA Hotairm1 was found to regulate granulocytic differentiation and HOX gene expression through a yet unknown mechanism (36, 37). During terminal myeloid cell differentiation, PU.1-induced lncRNA *lnc-MC* was reported to promote monocyte-to-macrophage differentiation (38). These seminal studies support the notion that lncRNAs critically contribute to the timing of myelopoietic programs and suggest that PIRAT is embedded into a larger regulatory RNA network in myeloid cells during homeostasis and disease.

Beyond its relevance to COVID-19, the specific expression of PIRAT in the myeloid lineage also renders this lincRNA a convenient biomarker of myeloid cell infiltration in bulk qRT-PCR or transcriptomics analysis of patient samples. Measurements with bronchoalveolar lavage fluid from control patients and patients with bronchopulmonary infection or idiopathic pulmonary fibrosis exemplify the utility of PIRAT in this context (Fig. 7). Besides their utility as biomarkers, noncoding RNAs, such as PIRAT, governing human immune-circuits, should be considered potential pharmacological targets. Recent successes in antisense-directed therapies (39) make lncRNA inhibitors seem feasible. The application of our recently introduced transfection-free approach for antisense-manipulation of myeloid RNA-circuits to PIRAT could, for instance, enable a more specific adjustment of transcription factor activity in the myeloid compartment, than possible with conventional, broad acting small molecule drugs (40). Besides PIRAT, further immune-regulatory lncRNAs could become relevant in this context. MaIL1 for instance supports type I IFN expression and LUCAT1 limits type I IFN immunity and STAT-dependent ISG expression in human macrophages (12, 25). Both lincRNAs are up-regulated during COVID-19 (Fig. 2G). STAT pathway inhibition by Ruxolitinib has been reported to prevent the progression of COVID-19 with systemic hyperinflammation into multiorgan-failure (41). Thus, whereas MaIL1 could nurture COVID-19 pathogenesis, LUCAT1 might adopt a protective function during COVID-19, preventing from excessive STAT-driven immune responses. The anticipated relevance of MaIL1 and LUCAT1 and further immune-regulatory lncRNAs, such as GAPLINC, PACER or CARLR to COVID-19 pathogenesis, and their utility as therapeutic targets, however, remains to be demonstrated.

In summary, our results suggest a multi-staged model of inflammation control in COVID-19 and other infectious diseases, in which lncRNAs occupy a central position. In the myeloid system, lncRNAs such as PIRAT, LUCAT1 and MaIL1 control the activity of immune master-transcription factors such as PU.1, STAT1 and IRF3 via complex feedback mechanisms. Whereas MaIL1 and LUCAT1 govern type I IFN immunity, negative feedback between PU.1 and PIRAT ensures that down-stream production of the critical alarmins S100A8/A9, involved in COVID-19 pathogenesis, is kept within narrow limits. Correspondingly, malfunctions at the lincRNA level can have a decisive influence on the course of COVID-19 and other immune-associated diseases. Further exploration of the nuclear RNA-layer that governs human myeloid immune responses will greatly improve our molecular understanding of the pathogenesis of infectious diseases and pave the way for more tailored and targeted interventions.

## Materials and Methods

### Cell culture and human biomaterial

Human peripheral blood mononuclear cells from healthy donors (control patient cells see below) were isolated from buffy coats (transfusion medicine department, UKGM Giessen). Buffy coats were de-identified prior to further use. Leukocyte populations were purified from buffy coats using Lymphoprep gradient medium (Stemcell Technologies) and MACS-purification (Miltenyi CD14-, CD4-, CD8-, CD45RO-, CD19-, CD56- and CD66b-beads). CD4 and CD8 T-cells were separated into CD45RO-positive and -negative populations, respectively. Blood-derived macrophages and dendritic cells were obtained by cultivating monocytes in the presence of 100 ng / ml GM-CSF or 50 ng / ml GM-CSF, 20 ng / ml IL-4 (Preprotech), respectively, in X-Vivo 15 medium (Lonza), containing 5 % fetal calf serum (FCS, Biochrom) for 7 days. Cell populations shown in Fig. 2H were isolated by cell sorting of gradient-purified leukocytes according to the following surface markers. Plasmacytoid DCs: CD19-, CD3-, CD56-, HLA-DR+, CD11C-, CD14-, CD16-, CD304+. Myeloid CDs: CD19-, CD3-, CD56-, HLA-DR+, CD11C+, CD14-, CD16-, CD1c/CD141+/-. Classical monocytes: CD19-, CD3-, CD56-, HLA-DR+, CD11C+, CD14+, CD16-. Non-classical monocytes: CD19-, CD3-, CD56-, HLA-DR+, CD11C+, CD14lo, CD16+. THP1 and Hek293T cells were purchased from ATCC and cultured in RPMI 1640 (Thermo Fisher), 10 % FCS, 1% penicillin/streptomycin solution (Thermo Fisher). For inhibitor treatments, cells were pre-stimulated with the respective inhibitors or DMSO for 2 h prior to further stimulations. Cells were cultured at a density of 1 million cells per 2 ml culture medium in 6-well dishes or with evenly adjusted cell number and medium volume for smaller dishes. In all experiments, LPS was used at a concentration of 100 ng / ml, polyI:C at 10 µg / ml and Pam3CSK4 at 200 ng / ml. All cells were cultured at 37 °C in a humidified atmosphere with 5 % CO_2._

Patients suffering from SARS-CoV-2-infection were recruited after hospitalization. In addition, healthy subjects were recruited (Table S3 and 4). All COVID-19 patients were tested positive for SARS-CoV-2 RNA in nasopharyngeal swabs and graded to have mild (WHO 2-4) or severe (5-7) disease according to the WHO clinical ordinal scale. Immunosuppressed, pregnant and HIV-positive patients were excluded from the study. The BioInflame study was approved by the ethics committee of the Charité - Universitätsmedizin Berlin (EA2/030/09) and the University Medical Center Marburg (55/17). All blood donors were at least 18 years of age and provided written informed consent for use of their blood samples for scientific purposes. PBMCs were isolated by Pancoll gradient centrifugation of one collected Vacutainer EDTA-tube (6 ml whole blood). All methods were performed in accordance with the relevant guidelines and regulations.

Bronchoalveolar lavage (BAL) fluid (BALF) (Fig. 7C) was obtained at the University Clinics Giessen and Marburg (UKGM) (American Thoracic Society consensus procedure), on approval by the ethics committee (Marburg: 87/12). Late-stage IPF tissue was analysed by the DZL/UGMLC biobank Giessen on approval by the ethics committee (AZ 58/15). Tissue was flushed with pre-warmed PBS. Obtained cells were analysed immediately. Further BALF (Fig. 7B) was obtained from patients at the Department of Infectious Diseases and Respiratory Medicine, Charité, Berlin. All patients underwent bronchoscopy including BAL on clinical indication and had provided oral and written informed consent. The study was approved by the local ethics committee (EA2/086/16). BAL was performed by instillation of 150 ml pre-warmed sterile 0.9% NaCl solution. In patients with focal abnormalities in chest imaging, BAL was performed in the corresponding pulmonary segment; in patients without radiological abnormalities or diffuse infiltrates, BAL was performed in the right middle lobe or lingula. Diagnosis of infection was made by a board-certified pulmonologist based on chest imaging, clinical signs of infection, culture and laboratory results, BALF cellular analysis and response to therapy. For the infection group, patients with non-mycobacterial infection were selected. Control patients showed no apparent lung disease and underwent bronchoscopy and BAL as part of rule-out diagnostics due to idiopathic coughing, for exclusion of pulmonary involvement of systemic disease (e.g. rheumatoid arthritis) or for exclusion of pulmonary tuberculosis. No obvious abnormalities in chest imaging and BALF composition were detected in these patients. Patient characteristics are listed in Table S5.

### CRISPR/Cas9

PIRAT-deficient cells were generated by CRISPR/Cas9, as recently described (42), using independent gRNAs (Fig. S6A). PIRAT gRNA pair -/-: GATGAGTCTAACGTGCACCC, GAGAGTTATAACATAATGGT. PIRAT gRNA pair +/-: ACGGATGGCCTTGGTCACCC, TTACATGAATAGACAGCTAG. Control cells were generated using a pX458 vector with scrambled gRNA. For PU.1 silencing, a lentiviral CRISPR interference vector (43) was used (Addgene #71237). gRNAs targeting the PU.1 TSS were cloned into the vector followed by lentiviral particle production (see below) and transduction; transduced cells (GFP+) were purified by cell sorting (Aria III, BD) and lysed immediately.

### Lentiviral transduction

HEK293T cells were co-transfected with lentiviral vector, pseudotyping- and helper-plasmid (pVSVG and psPAX2) using lipofectamine 2000 (Thermo Fisher). For over-expression, the SparQ lentivector (Systembio, # QM511B-1) containing the RACE-refined PIRAT cDNA was used. Virus-containing supernatants were passed through a 0.45 µm filter. Cells were transduced by resuspension in virus containing supernatants and centrifugation at 37 °C and 800 g for 2 h. 48 h later, transduced cells were purified by cell sorting (Aria III, BD) based on GFP-expression.

### qRT-PCR

RNA was isolated using TRIzol (Ambion), treated with DNaseI (Thermo Fisher) in the presence of recombinant RNase inhibitor (Promega) and concentration was determined (Nanodrop 2000 spectrometer, Thermo Scientific). cDNA was generated (High-Capacity cDNA Reverse Transcription Kit, Thermo Fisher) and quantitative PCR was performed (PowerUP SYBR Green Master Mix, Thermo Fisher) using a QuantStudio 3 instrument. For subcellular fractionation and CoIP analysis the Power SYBR RNA-to-Ct 1-Step Kit (Thermo Fisher) was used. Relative expression was calculated based on CT values, using the 2^-ΔΔCT^ method (44), where applicable relative to U6 snRNA. Primers are listed in Table S6.

### RACE-PCR

RACE-PCR was performed using the SMARTer 5’/3’ RACE kit (Clontech) according to the manufacturer’s instructions. Template poly(A) RNA was purified using oligo-d(T) coupled dynabeads (Thermo Fisher). RACE-PCR primers are listed in Table S6. RACE products were subjected to gel-purification and sub-cloned using the Strataclone UA PCR cloning kit (Agilent). Insert sequences were determined by Sanger sequencing (Seqlab GmbH).

### Subcellular fractionation

Cells were lysed (10 mM Tris, pH 8, 140 mM NaCl, 1.5 mM MgCl_2_, 0.5 % Igepal, 2 mM vanadyl ribonucleoside complex), incubated on ice for 5 min and centrifuged (1000 x g, 4 °C, 3 min). The supernatant (cytosolic fraction) was transferred to a new tube, centrifuged (3 min, maximum speed) and transferred to a new tube for RNA-extraction. The pellet (nuclear fraction) was washed two times with lysis buffer and once with lysis buffer containing 0.5 % deoxycholic acid (centrifugations at 4 °C and 1000 x g), followed by RNA-extraction.

### RNA-FISH

Tissues were derived as described (12) with Charité University Medicine Berlin Ethics Committee approval no. EA2/079/13, formalin fixed, paraffin embedded and sectioned at 4 µm on glass slides. Probe sequences were designed by Affymetrix (Homo sapiens PIRAT (RUO) Catalog no. VA1-3025697; Homo sapiens EEF1A1 Catalog no. VA1-10418). RNA-FISH was performed using the ViewRNATM ISH Tissue 1-Plex Assay (Affymetrix) with heat pretreatment for 10 min and protease digestion for 20 min. A probe homologous to EF1α served as positive control for the hybridization conditions on consecutive tissue sections. Diluent without probe served as control for background staining. Roti®-Mount FluorCare DAPI (Carl Roth®) was used for counterstaining of nuclei and as mounting medium. Photographs were taken using an Olympus DP 80 microscope at 600x magnification (DAPI signal: 345 nm; red probe signals: 550 nm).

### Western blot

Protein concentrations were determined using BCA (Pierce™ BCA Protein Assay Kit, ThermoFisher) and an Infinite PRO (Tecan) plate reader. Proteins were separated by SDS PAGE, using 10% polyacrylamide gels. Proteins were transferred onto a nitrocellulose membrane (Amersham™ Protran®, Sigma-Aldrich). For blot development and detection, the ECL Prime Western Blot Detection kit (Amersham) and a Chemostar Imager (INTAS Science Imaging) were used. Antibodies are listed in Table S8. Western blot full-scan is shown in Fig. S7B.

### Flow cytometry

Cells were identified by plotting the respective fluorescence channel against background-fluorescence or the side-scatter. The gating strategy is illustrated in Fig. S6D. For surface marker staining, 2 µl of fluorophore-coupled primary antibody were added to cells in 100 µl PBS containing 1 % FCS, followed by incubation on ice for 30 min. Cells were washed and resuspended in PBS containing 0.5 % FCS and subjected to FACS analysis (Guava EasyCyte, Millipore).

### ChIP

40 million cells per capture were crosslinked with PBS, 1 % formaldehyde for 10 min, quenched with 1/10th volume 1.25 M glycine for 5 min and resuspended in 800 µl lysis buffer (50 mM Tris-Cl, 10 mM EDTA, 1 % SDS, 1 mM PMSF). Lysate was sonicated (Diagenode Biorupter) until DNA appeared with a fragment size between 100 and 500 bp on agarose gels. Sample was adjusted with 3.6 ml ChIP Dilution Buffer (50 mM Tris-HCl, 0.167 M NaCl, 1.1% Triton X-100, 0.11% sodium deoxycholate), 2 ml RIPA-150 (50 mM Tris-HCl, 0.15 M NaCl, 1 mM EDTA pH8, 0.1% SDS, 1% Triton X-100, 0.1% sodium deoxycholate) and PMSF (1 mM). 60 µl of magnetic beads were coupled with PU.1 C1 + A7 antibody or FLAG antibody (Table S8), as described by Tawk et al. (45) and added to the diluted lysate, followed by rotation at 4 °C over-night. Upon one wash with RIPA-150, two washes with RIPA-500 (same as RIPA-150 but with 0.5 M NaCl), 2 washes with RIPA-LiCl (50 mM Tris-HCl, 1 mM EDTA pH 8, 1% Nonidet P-40, 0.7% sodium deoxycholate, 0.5 M LiCl2) and 2 washes with TE buffer (10 mM Tris-HCl, 1 mM EDTA), DNA input and bead samples were resuspended in 200 µl elution buffer (10 mM Tris-HCl, 0.3 M NaCl, 5 mM EDTA pH8, 0.5 % SDS). Until this step, all buffers were supplemented with cOmplete protease inhibitor (Roche). Following addition of 1 µl RNase A and incubation for 4 h at 65 °C beads were separated and supernatant was incubated with 10 µl of Proteinase K for 45 min at 50 °C. DNA was purified by PCI extraction and ethanol / sodium-acetate precipitation.

### ChIRP

Antisense DNA probes (Table S7) were synthesized at Metabion AG and 3’ mono-biotinylated using terminal transferse (New England Biolabs) and Biotin-11-ddUTP (Jena Bioscience) according to the manufacturer’s instructions. ChIRP (20 million CD14+ monocytes per capture) was performed as described previously (46).

### UV crosslinking & Co-immunoprecipitation

For co-immunoprecipitation (CoIP), 10^7^ cells were UV-crosslinked (300 mJ / cm^2^) in petri dishes, on an ice bath. The CoIP procedure published by Tawk et al. (45) was used with minor modifications. For protein purification protein G dynabeads (Thermo Fisher), coupled with 2.5 µg of antibody (Table S8) were used. In PU.1 CLIP experiments, eluate fractions were split up for protein analysis by Western blot and RNA extraction as described above.

### Single Cell RNA-sequencing analysis

Single cell multiomics was performed using the BD Rhapsody system according to manufacturer’s protocols. 250.000 PBMCs per sample (two patients and two healthy controls) were incubated with an individual oligo-labelled antibody (Multiplex Tag, BD Human Single-Cell Multiplexing Kit, Cat. No. 633781), for 20 minutes at room temperature. Cells were washed twice with BD Pharmingen Stain Buffer (Cat. No. 554656) and labelled cell suspensions were pooled and incubated with oligo-labelled AbSeq antibodies directed against CD206 (Cat. No. 940068), CD163 (Cat. No. 940058) and HLA-DR (Cat. No. 940010) for 30 minutes on ice. Upon two washes with Stain Buffer, cells were resuspended in Sample Buffer (Cat. No. 650000062) and viability-stained with 2 mM Calcein AM (Cat. No. C1430; Thermo Fisher Scientific, Dreieich, Germany) and 0.3 mM Draq7 (Cat. No. 564904) for 5 minutes at 37°C. The suspension was counted using a disposable hemocytometer (Cat. No. DHCN01-5; INCYTO, Cheonan, South Korea) and cell viability was determined.

The BD Rhapsody Cartridge (Cat. No. 400000847) was primed with 100% ethanol followed by 2 washes with Cartridge Wash Buffer 1 (Cat. No. 650000060) and one wash with Cartridge Wash Buffer 2 (Cat. No. 650000061). About 30.000 labelled cells were loaded and incubated for 15 minutes at room temperature. Excess fluid was removed and the cartridge was loaded with Cell Capture Beads (Cat. No. 650000089) and incubated for 3 minutes at room temperature. Excess beads were washed off using Sample Buffer. Lysis Buffer was applied and beads were extracted from the cartridge using the BD Rhapsody Express instrument and washed twice with cold Bead Wash Buffer (Cat. No. 650000065).

The cDNA reaction mix was prepared as indicated in the manufacturer’s protocol and mixed with the beads. The mixture was incubated in a thermomixer (37 °C, 1200 rpm, 20 minutes). The supernatant was removed and replaced by the Exonuclease I mix prepared according to the manufacturer’s protocol. The bead suspension was placed on the thermomixer (37°C, 1200 rpm, 30 minutes, followed by 80 °C without shaking for 20 minutes). The suspension was then briefly placed on ice and the supernatant was removed. Finally, beads were resuspended in Bead Resuspension Buffer (Cat. No. 650000066).

Single cell mRNA, multiplex sample Tag, and AbSeq libraries were prepared using the BD Rhapsody™ Single-Cell Analysis system (Cat. No. 633774) according to manufacturer’s recommendation (Doc ID: 214508). Briefly, the Bead Resuspension Buffer was removed from the beads and replaced by the PCR1 reaction mix containing the primers specific for the AbSeq and mutiplex sample tags, and genes of the Human Immune Response Panel (Cat. No. 633750) supplemented with custom-made primers for additional genes (see NCBI GEO GSE142503). Beads were placed in the thermal cycler for 11 cycles of the PCR program indicated in the protocol. The supernatant was retained and the PCR products for Abseq and multiplex sample tags, as well as the mRNA PCR product were separated and purified by double-sided size selection using AMPure XP magnetic beads (Cat. No. A63880; Beckman Coulter, Krefeld, Germany). A fraction of the Abseq/multiplex sample tag PCR 1 product, as well as the mRNA PCR 1 product were further amplified with a second PCR of 10 cycles and subsequent purification using AMPure XP beads, resulting in multiplex sample tag and mRNA PCR 2 product. Finally, the Abseq/multiplex sample tag PCR 1 product for the Abseq library, and both PCR 2 products for each the multiplex sample tag and mRNA libraries were amplified by the final index PCR for 7 cycles each with subsequent purification afterwards. Concentrations of the index PCR products were determined using the Qubit Fluorometer and the Qubit dsDNA HS Kit (Cat. No. Q32851; Thermo Fisher Scientific) and quality control was performed on the Agilent 2100 Bioanalyzer with the High Sensitivity DNA Kit (Cat. No. 5067-4626; Agilent, Waldbronn, Germany). Mixed libraries were sequenced on a NextSeq550 with 2 × 75 bp paired-end reads.

After pre-processing of BD Rhapsody scRNA-seq data, read counts were loaded into the R (v3.6.3) environment and further analyzed using the Seurat package (v3.1.4). The following quality criteria were used to include cells for the downstream analysis: at least 25 genes were expressed, and at least 1,000 but no more than 70,000 transcripts were detected per cell.

Following the Seurat workflow, the read counts were normalized and scaled by NormalizeData and ScaleData functions of the Seurat package, respectively. Principal component analysis (PCA) was performed by RunPCA using top 2,000 variable features that were selected using the default selection method (“vst”) in Seurat. Next, based on the first 15 PCs, cell clusters were identified with the Louvain algorithm at resolution of 0.4. Finally, in a two-dimensional space, a UMAP was generated to visualize the identified cell clusters.

To identify marker genes of each cell cluster, differentially expressed (DE) genes were tested by FindAllMarkers functions in Seurat using the default test (Wilcoxon Rank Sum test). Significantly differentially expressed genes were determined by 1) log-fold changes > 0.3, 2) expressed in at least a fraction of 0.2 cells in each tested population, and 3) adjusted p value < 0.05 (Bonferroni correction). DE analyses were used to identify cluster marker genes by comparing the expression of upregulated genes in cells between one cluster and the rest of cells.

Cell clusters were firstly assigned using the SingleR (v1.0.6) package based on four reference dataset which are provided in the package, including BlueprintEncodeData, DatabaseImmuneCellExpressionData, HumanPrimaryCellAtlasData, and MonacoImmuneData. Then, the assigned cell cluster annotations were double-checked by comparing the cell type specifically expressed marker genes from public resources.

To dissect the different profiles between COVID-19 patients and controls, publicly reported COVID-19 related genes were selected and their expression profiles in patients and controls were visualized for each identified cell cluster using a modified DotPlot function in Seurat.

### Bulk sequencing and bioinformatics analysis

RNA was isolated (miRVana kit, Thermo Fisher) and DNaseI-digested as described above. RNA-quality was evaluated (Experion RNA analysis kit, BioRad) and Illumina TruSeq mRNA libraries were generated (Genomics Core Facility, Philipps-University Marburg), and analysed on a HiSeq 1500 machine. CLIP-seq and ChIRP-seq libraries were generated at Vertis Biotech AG (Germany) using in-house protocols and sequenced on a NexSeq500 device. Human Bodymap raw data (Fig. S1A) were obtained through NCBI Sequence Read Archive (datasets ERR030888-ERR030903). Peripheral blood leukocyte raw data (Fig. 1B) were downloaded from NCBI GEO (GSE62408 and GSE60424). ENCODE CD14^+^-monocyte DNaseI-Seq, H3K4me3- and H3K27me3-ChIP-Seq data were downloaded from NCBI GEO (SRR608865, SRR608866, SRR568364, SRR568365, SRR568417, SRR568418). NCBI data were extracted using the SRA toolkit. Haematopoietic lineage expression raw data were obtained through the Blueprint Consortium (EGAD00001000939, EGAD00001000919, EGAD00001000907, EGAD00001000922, EGAD000010001477, EGAD00001000675).

Reads in fastq-format were quality-trimmed and mapped to the human GRCh38 reference (GENCODE), using the CLC genomics workbench. Gene expression changes were calculated using RPKMs (based on uniquely mapped reads). Hierarchical clustering was done using Cluster 3.0 (Eisen lab). Heatmaps were generated using JAVA TreeView (47). For pathway enrichment analysis and induced network analysis ConsensusPathDB (48) was used. For co-expression analysis R^2^ was calculated (Excel) based on RPKMs, and ENSEMBL-IDs of genes with R^2^ values ≥ 0.8 were analysed in ConsensusPathDB. PCA analysis was done based on row Z-scores, using the R-script prcomp (stats) with rgl package. Other plots were generated using GraphPad Prism, Excel or BoxPlotR (http://shiny.chemgrid.org/boxplotr/). Statistical analysis were performed using GraphPad Prism. Sequence conservation was determined using NCBI BLASTN and the major species reference genome, respectively. BLAST hits with ≥ 20 complementary nucleotides located within a genomic range of max. 100 kb were considered.

### Statistical analysis

Throughout this study, statistical analysis were performed based on ≥ three independent experiments, except for single cell RNA-seq experiments. Test details can be found in the figure legends and methods details. If not specified differently, GraphPad Prism software was used for two-tailed Student’s t-test and ANOVA analysis. Differences between two or more compared conditions were regarded significant when p-values were ≤ 0.05. Where possible, p-values are shown in the respective figure panels.

## Acknowledgements

We would like to thank Kerstin Hoffmann for assisting in cell culture experiments and Kathleen Stabla and Jennifer Kremer for assisting in patient sample collection. This study makes use of data generated by the Blueprint Consortium. A full list of the investigators who contributed to the generation of the data is available from www.blueprint-epigenome.eu. Funding for the project was provided by the European Union’s Seventh Framework Programme (FP7/2007-2013) under grant agreement no 282510 – BLUEPRINT.

This work was funded by the Deutsche Forschungsgemeinschaft SFB-TR84 “innate immunity of the lung” (project C10, to LNS and LES; project C1, to BS; project Z1b, to ADG), the Hessian Ministry of Higher Education, Research, Science and the Arts (LOEWE Diffusible Signals, to LNS and BS) and von Behring Röntgen Stiftung (project 63-0036, to LNS). SMV was supported by the Jürgen Manchot Foundation (Doctoral Research Fellowship). YL was supported by an ERC Starting Grant (948207) and the Radboud University Medical Centre Hypatia Grant (2018) for Scientific Research.

## Author contributions

Experimental work: MA, NS, HJ, ZZ, KP, JH, SMV, DW, PG, MG, EM, CS, CK, CB, WB, AN, TS, ADG, CR, YL, HG, LES, BS, LNS

Manuscript writing and figure preparation: LNS, MA, ZZ, YL, JH

Proof-reading: MA, NS, HJ, ZZ, KP, JH, SMV, DW, PG, MG, AN, TS, ADG, CK, CR, YL, HG, LES, BS, LNS

Project supervision: LNS

## Data availability

Bulk and single-cell RNA-seq data have been deposited at NCBI GEO (GSE142503) and are publicly available as of the date of publication. All other data will be shared by the corresponding author upon request.

## Supplementary Information

Figures S1 to S7

Tables S1 to S8

