## Supplementary Information for "Single cell RNA-seq uncovers the nuclear decoy lincRNA PIRAT as a regulator of systemic monocyte immunity during COVID-19"

Fig. S1. Identification of myeloid and lymphoid lincRNAs signatures.
Fig. S2. PIRAT co-expression network.
Fig. S3. PBMC single cell RNA-seq data analysis.
Fig. S4. Monocyte response in COVID-19 (differential gene expression).
Fig. S5. Characterization of PIRAT cDNA sequence by RACE PCR
Fig. S6. Basic properties of lincRNA PIRAT.
Fig. S7. Characterization of PIRAT interplay with PU.1.

Table S1. Myeloid and lymphoid cell specific lincRNAs identified in the current study.
Table S2. List of PU.1 target genes.
Table S3. COVID patient characteristics (cohort 1).
Table S4. COVID patient characteristics (cohort 2).
Table S5. Characteristics of control and pulmonary infection patients.
Table S6. PCR and sequencing oligonucleotides used in the present study.
Table S7. ChIRP oligonucleotides used in the present study.
Table S8. Antibodies used in the present study.

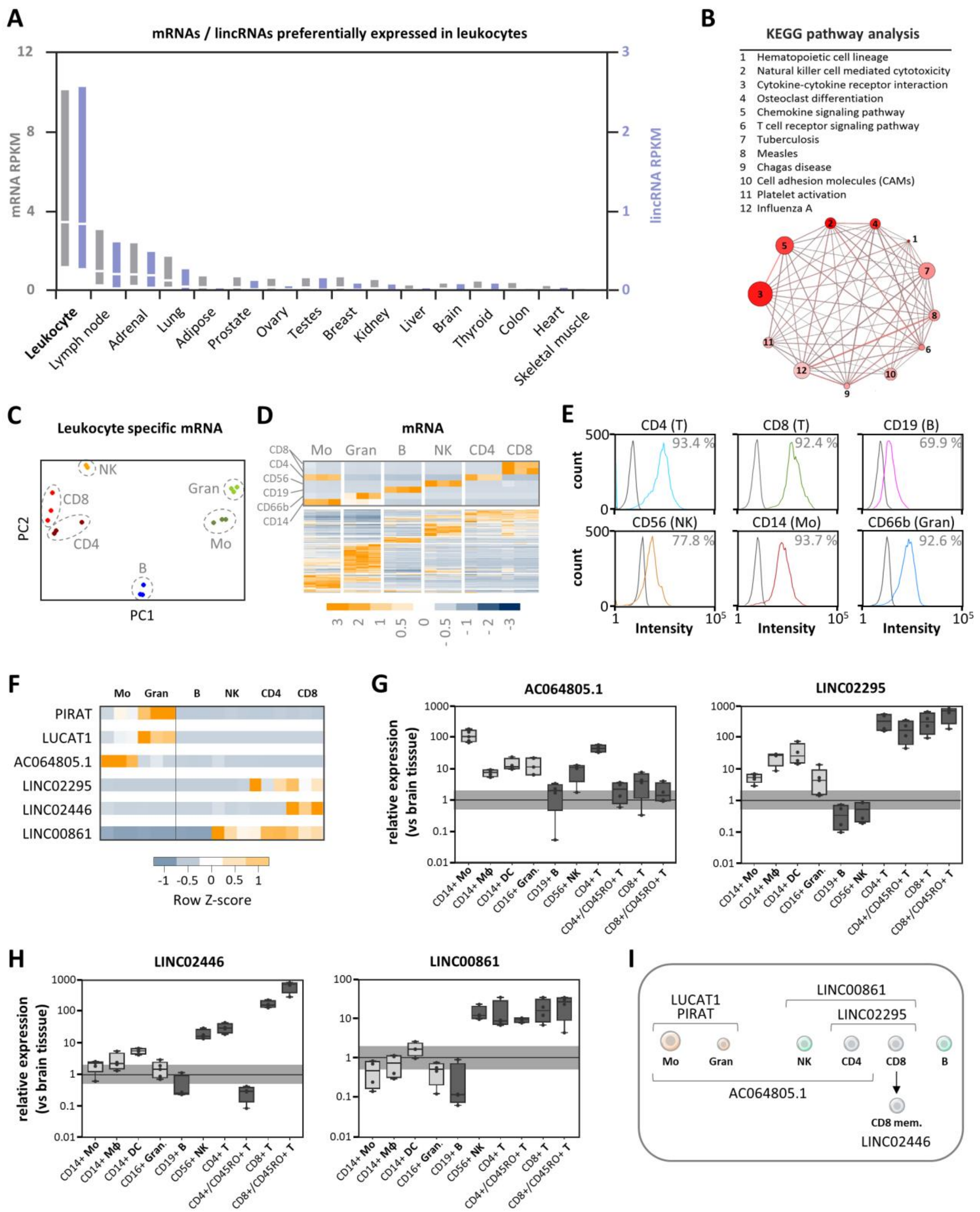

**Fig. S1. Identification of myeloid and lymphoid lincRNAs signatures.** A) mRNA and lincRNA expression levels (median + inner quartiles) in indicated tissues (Human Bodymap). B) KEGG pathway analysis of mRNA from A. Pathway size = circle size. Candidates contained: color-coded (light to dark red). Overlap between pathways = line thickness. C) and D) PCA analysis with leukocyte-enriched lincRNAs and clustering of mRNAs from A (cell type markers indicated). E) Overlay histogram FACS plots showing successful enrichment of the indicated cell populations by MACS (left peak: unstained control; right peak: cells stained for indicated surface antigen). F) RNA-Seq based row Z-scores of selected myeloid (top 3)

and lymphoid (bottom 3) lincRNA markers (data from Fig. 1B). G) and H) qRT-PCR validation of lincRNAs from indicated purified cell types, relative to human brain reference tissue. Horizontal bar indicates base-line (black) and 2-fold deviation from base-line (grey). Box plots and individual replicate values from four independent experiments are shown. I) Summary of lincRNA expression patterns in the studied leukocyte populations.

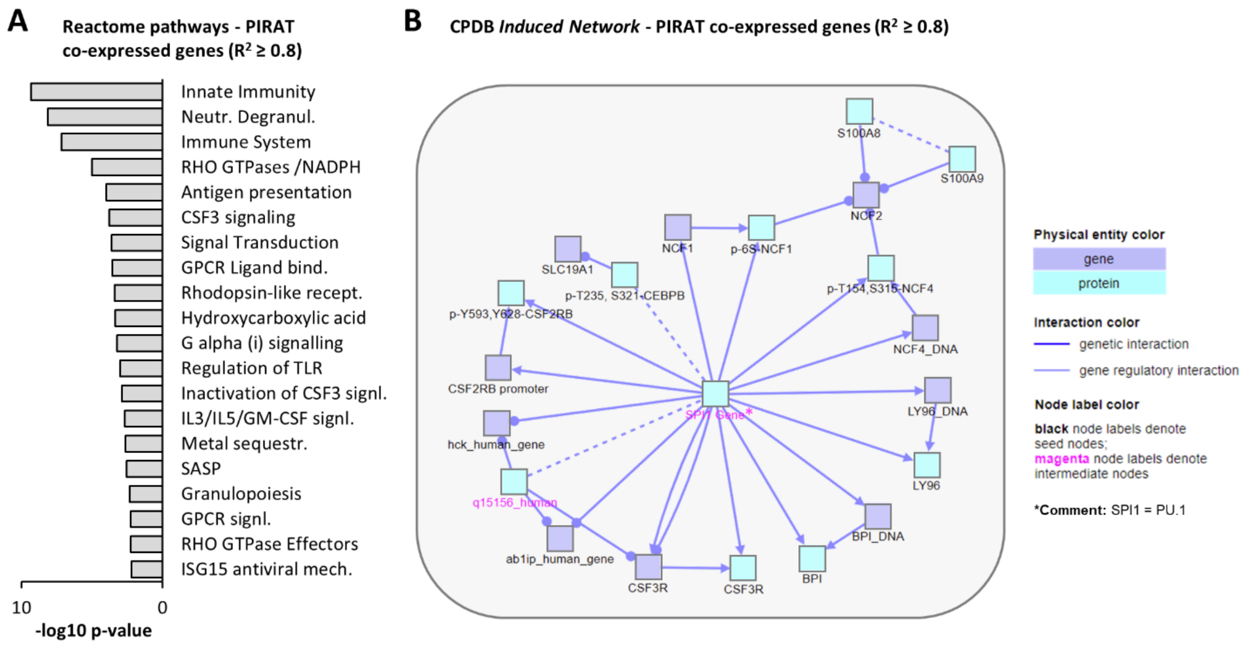


**Fig. S2. PIRAT co-expression network.** A) Consensus Path DB reactome pathway analysis and B) Consensus Path DB induced network analysis of genes co-expressed with PIRAT in RNA-seq datasets from Fig. 1.

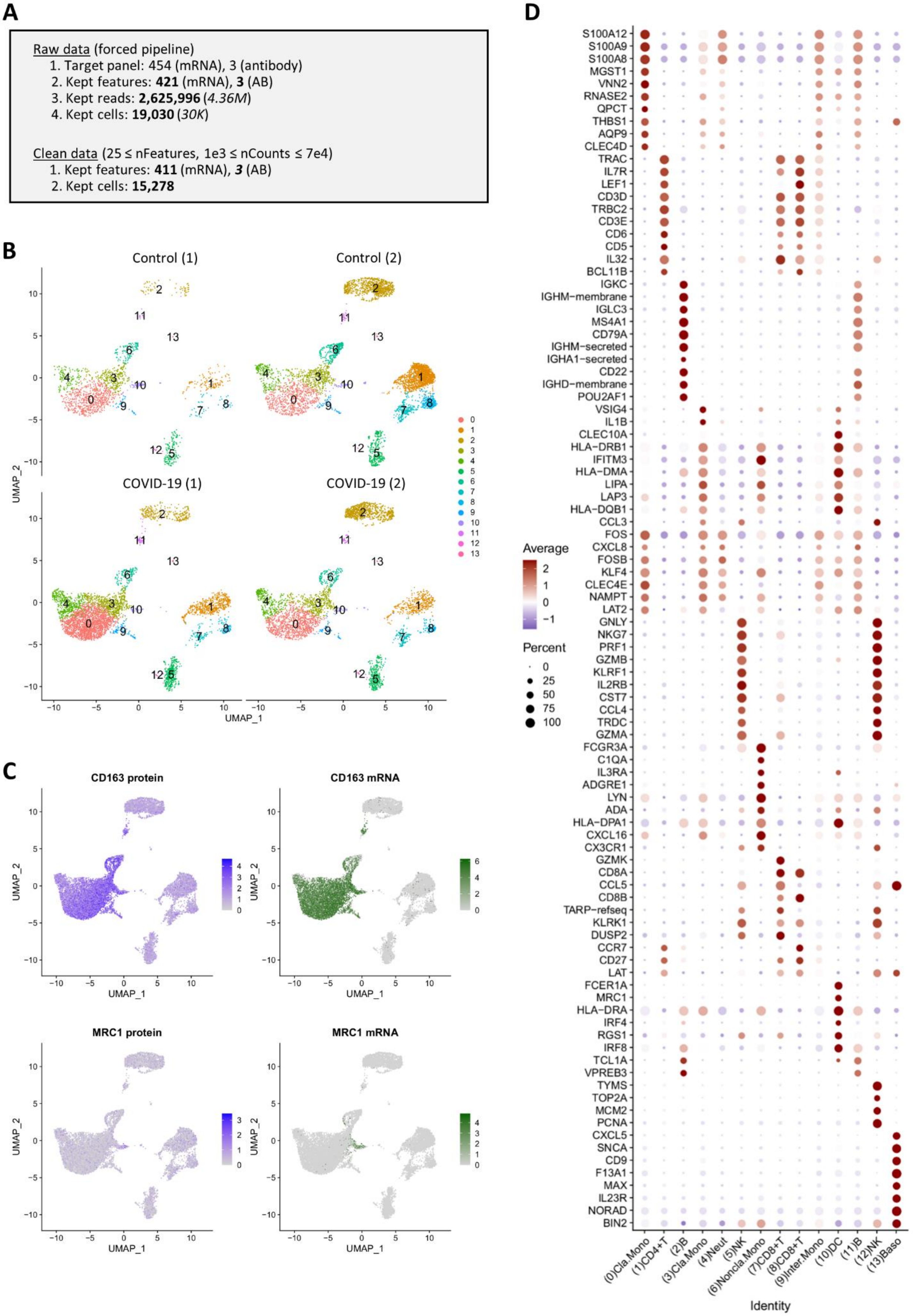
A
Raw data (forced pipeline)
1. Target panel: 454 (mRNA), 3 (antibody)
2. Kept features: 421 (mRNA), 3 (AB)
3. Kept reads: 2,625,996 (4.36M)
4. Kept cells: 19,030 (30K)
Clean data (25 ≤ nFeatures, 1e3 ≤ nCounts ≤ 7e4)
1. Kept features: 411 (mRNA), 3 (AB)
2. Kept cells: 15,278
B
Control (1)
Control (2)
COVID-19 (1)
COVID-19 (2)
UMAP_1
UMAP_2
C
CD163 protein
CD163 mRNA
MRC1 protein
MRC1 mRNA
D
S100A12
S100A9
S100A8
MGST1
VNN2
RNASE2
QPCT
THBS1
AQP9
CLEC4D
TRAC
IL7R
LEF1
CD3D
TRBC2
CD3E
CD6
CD5
IL32
BCL11B
IGKC
IGHM−membrane
IGLC3
MS4A1
CD79A
IGHM−secreted
IGHA1−secreted
CD22
IGHD−membrane
POU2AF1
VSIG4
IL1B
CLEC10A
HLA−DRB1
IFITM3
HLA−DMA
LIPA
LAP3
HLA−DQB1
CCL3
FOS
CXCL8
FOSB
KLF4
CLEC4E
NAMPT
LAT2
GNLY
NKG7
PRF1
GZMB
KLRF1
IL2RB
CST7
CCL4
TRDC
GZMA
FCGR3A
C1QA
IL3RA
ADGRE1
LYN
ADA
HLA−DPA1
CXCL16
CX3CR1
GZMK
CD8A
CCL5
CD8B
TARP−refseq
KLRK1
DUSP2
CCR7
CD27
LAT
FCER1A
MRC1
HLA−DRA
IRF4
RGS1
IRF8
TCL1A
VPREB3
TYMS
TOP2A
MCM2
PCNA
CXCL5
SNCA
CD9
F13A1
MAX
IL23R
NORAD
BIN2
Average
Percent
(0)Cla.Mono
(1)CD4+T
(2)B
(3)Cla.Mono
(4)Neut
(5)NK
(6)Noncla.Mono
(7)CD8+T
(8)CD8+T
(9)Inter.Mono
(10)DC
(11)B
(12)NK
(13)Baso
Identity

**Fig. S3. PBMC single cell RNA-seq data analysis.** A) Illustration of data analysis strategy and statistics. B) UMAP plots showing the clustering of cell types in the indicated control and COVID-19 patient scRNA-seq datasets. C) UMAP plots showing surface protein marker (AbSeq) and mRNA detection for CD163 (myeloid marker) and MRC1 (dendritic cell / monocyte marker) in aggregated control and COVID-19 patient scRNA-seq data. D) Dot plot showing the color-coded average expression and the percentage of positive cells for characteristic mRNA/lincRNA markers in the indicated cell populations.

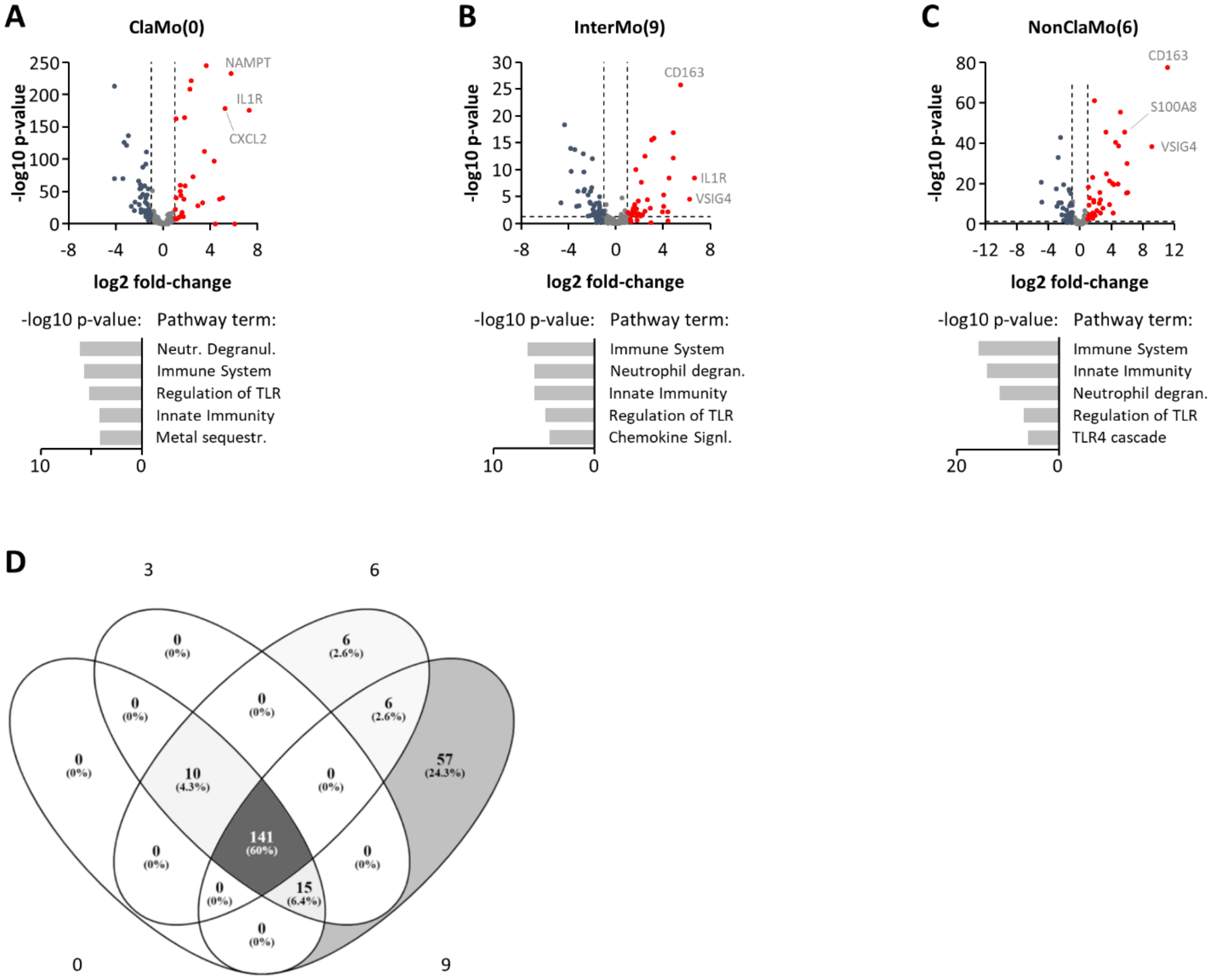


**Fig. S4. Monocyte response in COVID-19 (differential gene expression).** A) to C) Top: volcano plots showing gene expression changes in classical, intermediate and non-classical monocyte populations in COVID-19 compared to control patients (based on scRNA-seq data shown in Fig. S3). Top 3 induced mRNAs are indicated. Bottom: Consensus Path DB Reactome pathway analysis with significantly up-regulated mRNAs (top 5 pathways are shown). D) Overlap of differentially expressed genes in the respective monocyte populations (numbers denote cell populations defined in Fig. 2C and Fig. S3D).

### A 5'/3' RACE-PCR result

**>PIRAT_cDNA_sequence**

```
GAGGAACAGTCTTACTCTGTCACCCAGGCTGCAGTGTAGTGGTGTGATCACAGCTCACTGCAGCCTTGACCTCCTGG
GCTTAGGTGATCCTCCCACCCTAGCCTCCCATGTAGCTGGGACTAGAGGTATGTGCCACCTCACCTCCTTTTTTTCT
TTTCTTTTTCTTTTTTGGAGAGACAGATTCTTCTTATGTTGCTATTTTAAACTCCTGAACTCAAGTGATCCTCCTGC
CTTGGCCTCCCAAAGTGCTGGGATTACAGGTGTGAGACACTGCACCCTGCCCAAGCACCTCTGCTCCGTGCCATGCT
CTTGGCTAATTGGAGTTGTGAAAGGCATGAGGATTCTGGTCATGGACTCAGCTCTCCCATAGGGTTCTGACACCAAA
GCAATGGTCACAACAGTGAAAGGAAGAATCCATCTGGCCAGGCTCAGGTGGTTCAAGGCCTTCAGAATCTGCCTTGA
GACTCTCACTGGCTTTAGACTGAAAACCATCTTGGCCCCGTCCATCCGTGTAAGCAATTTAACGACAGCTTGCAAAG
CACCGAGCTTTAACAGAAAGAAGAGATGAGCACAGCGCAAGAACTTGGACTCCAGAAGAGCTGCCTAACAGATTATT
TTTCTGTGGCATTTCATGAGAACAAACGAAGTAGGAATTTTCCTTTTGTTTGTCTGGCCTTTGGCATCGTTTACTTT
CTTTTTATTCTTCTGAAATGTACTTCGAGCCCTGGCAGCATTTCTGTCCTAAAATCTTATTGTCAGAGGTTTATTTT
TCAGCTTTTCAAATCATATCTGATAGAGTGAGTGTACTGCCTGGACTCATCACTTTACTTCAGAAGAAATACAGCTC
ACCCTTTAAATGACAATGGTGACTGTCCACATCTTTATGTTTTCTACACTGAAGTGGCAGGCTTCATTTAAAAATAA
TGTTTTCCCTCATCAAAAGAGAGCTAGGGTAGAACCGTCAACTCTGCTGTTGTCTGGGTAGTGACCTAACACCCACG
TTTTGGACAATCACTCACTGTCTTATATTGGGTTTTCATTGCATGTAGGATAATTCTTTGTCAATGGTAGTTTTGTC
AACCGTGATCTGAGGTAATGAGGTTTTCTACTTTTGCTTGAAATTTTGAAAATATGCAAGCTTTAAACATTT
```

**Fig. S5. Characterization of PIRAT cDNA sequence by RACE PCR.** A) Full-length PIRAT sequence in human monocytes, reconstructed from Rapid Amplification of cDNA Ends (5' and 3') experiments and subsequent full-length PCR amplification and Sanger sequencing.

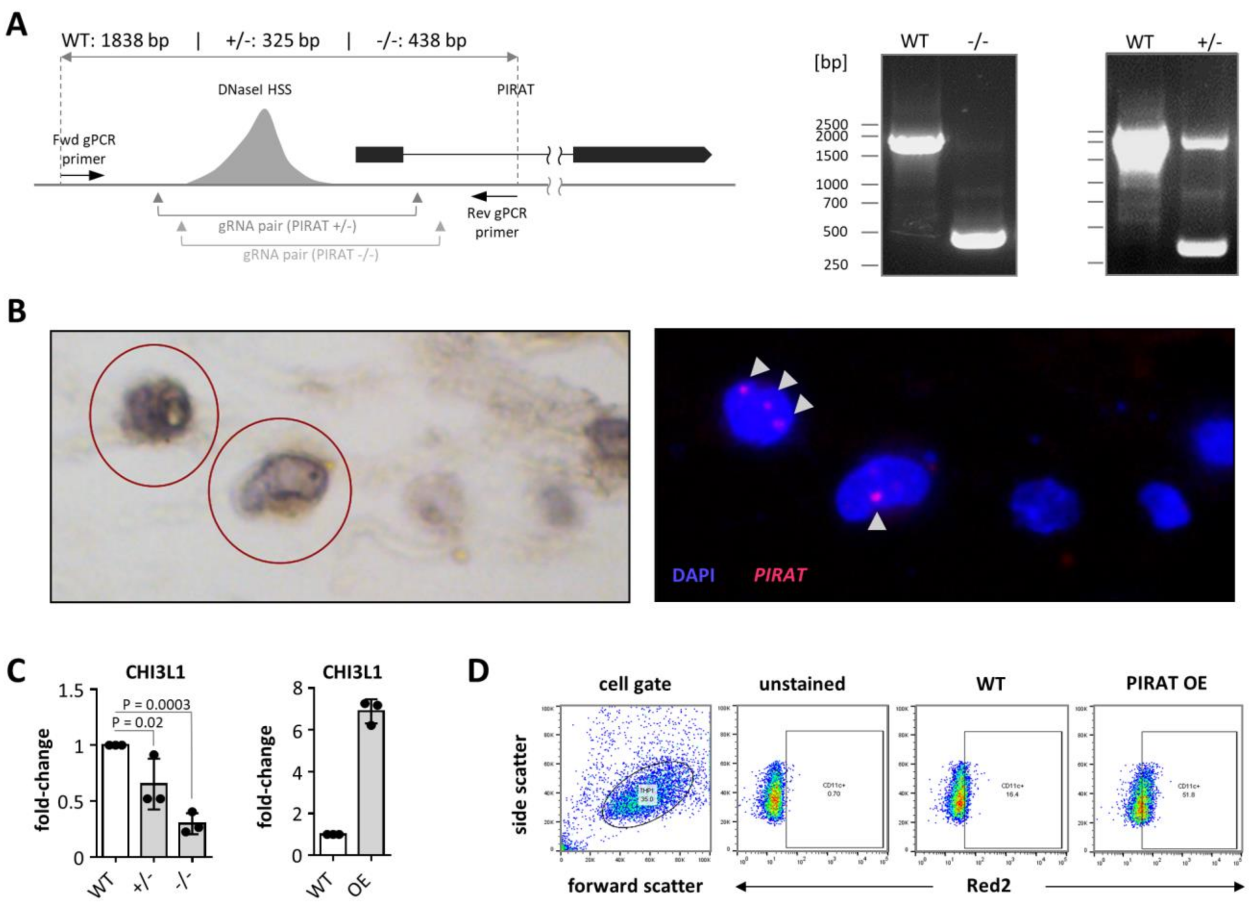


**Fig. S6. Basic properties of lincRNA PIRAT.** A) Left: schematic representation of guideRNA and genomic PCR primer binding sites in the PIRAT (LINC00211) locus. Expected Genomic PCR amplicon sizes for wild-type (WT), monoallelic (+/-) and biallelic knockouts (-/-) are indicated on the top. Right: Agarose gels showing genomic PCR amplicons from wild-type and PIRAT -/- and +/- THP1 cells. B) Left: hematoxylin staining of human lung slice. Circles indicate alveolar phagocytes. Right: RNA-FISH analysis of PIRAT subcellular localization in the same image. Nuclei counter-stained with DAPI. White arrows indicate PIRAT signal. C) qRT-PCR analysis of CHI3L1 expression in wild-type and PIRAT-deficient (+/- and -/-), as well as in PIRAT over-expressing (OE) THP1 cells. Mean, individual replicate values and standard deviation based on three independent experiments are shown. D) Representative dot plots illustrating the gating strategy used for FACS-quantification of CD11c (ITGAX) positive THP1 cells (Fig. 4F).

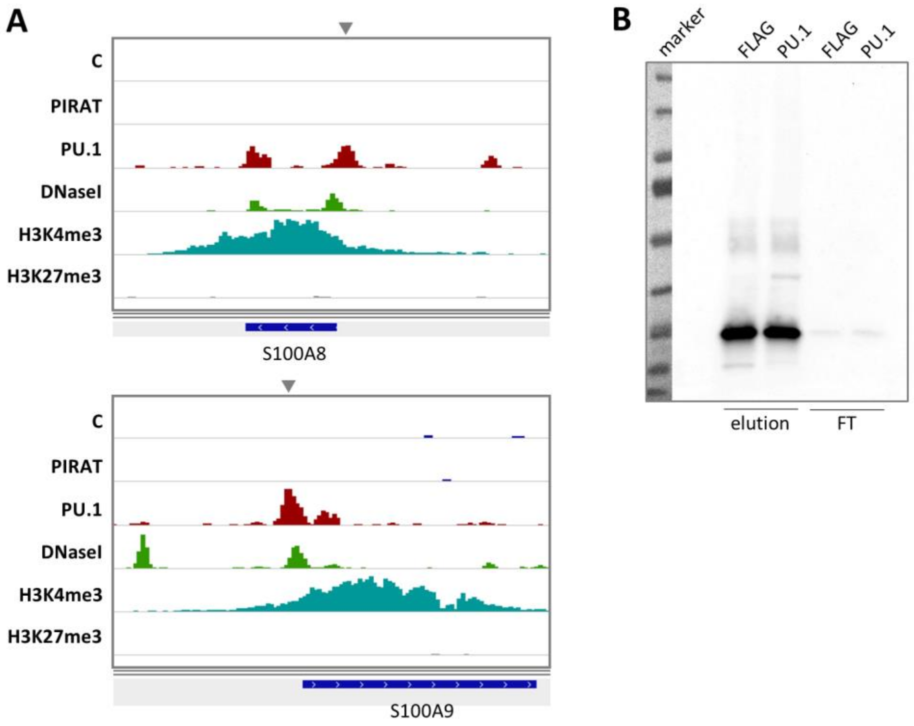


**Fig. S7. Characterization of PIRAT interplay with PU.1.** A) IGV plots showing PIRAT ChIRP-seq, PU.1 and histone H3 ChIP-seq and DNaseI-seq coverage in the S100A8 and A9 loci. Grey triangles indicate the assumed PU.1 proximal promoter binding sites. B) Full-scan of PU.1 CoIP Western blot shown in Fig. 6A (FT = flow-through fractions).

| Alias | Monocytes | Granulocytes | B cells | NK cells | CD4 T cells | CD8 T cells |
|---|---|---|---|---|---|---|
| AC064805.1 | 4.58 ± 1.19 | 0.23 ± 0.20 | 0.02 ± 0.03 | 0.09 ± 0.13 | 0.01 ± 0.01 | 0.00 ± 0.00 |
| AP001257.1 | 4.03 ± 1.73 | 0.01 ± 0.02 | 0.02 ± 0.04 | 0.02 ± 0.03 | 0.01 ± 0.02 | 0.03 ± 0.03 |
| AP003774.3 | 3.92 ± 1.83 | 0.06 ± 0.10 | 0.02 ± 0.04 | 0.04 ± 0.07 | 0.01 ± 0.01 | 0.01 ± 0.01 |
| LINC02285 | 3.69 ± 0.65 | 2.11 ± 1.06 | 1.20 ± 0.44 | 1.62 ± 0.56 | 0.92 ± 0.18 | 1.25 ± 0.19 |
| AC097504.2 | 3.53 ± 0.45 | 0.00 ± 0.00 | 0.00 ± 0.00 | 0.00 ± 0.00 | 0.00 ± 0.00 | 0.00 ± 0.00 |
| LUCAT1 | 2.49 ± 1.62 | 23.97 ± 15.61 | 0.02 ± 0.01 | 0.03 ± 0.02 | 0.02 ± 0.02 | 0.03 ± 0.03 |
| LINC01506 | 0.57 ± 0.09 | 9.27 ± 2.93 | 0.03 ± 0.01 | 0.00 ± 0.00 | 0.00 ± 0.00 | 0.00 ± 0.00 |
| AC007342.5 | 0.04 ± 0.05 | 8.38 ± 1.80 | 0.01 ± 0.02 | 0.00 ± 0.00 | 1.18 ± 0.67 | 1.27 ± 0.66 |
| PIRAT | 0.71 ± 0.47 | 4.63 ± 1.46 | 0.00 ± 0.00 | 0.04 ± 0.04 | 0.02 ± 0.03 | 0.02 ± 0.03 |
| LINC00921 | 0.52 ± 0.24 | 3.96 ± 1.01 | 0.14 ± 0.15 | 1.24 ± 1.30 | 0.33 ± 0.26 | 0.30 ± 0.29 |
| AC104971.4 | 0.06 ± 0.11 | 0.00 ± 0.00 | 7.52 ± 0.22 | 0.04 ± 0.04 | 0.50 ± 0.20 | 0.23 ± 0.20 |
| AC009686.2 | 0.00 ± 0.00 | 0.00 ± 0.00 | 5.18 ± 1.64 | 0.16 ± 0.28 | 0.16 ± 0.15 | 0.37 ± 0.30 |
| AC006033.2 | 1.17 ± 0.11 | 0.10 ± 0.01 | 0.02 ± 0.04 | 9.40 ± 1.20 | 0.02 ± 0.02 | 0.63 ± 0.22 |
| AC093323.2 | 0.16 ± 0.25 | 0.17 ± 0.10 | 0.06 ± 0.04 | 6.48 ± 1.97 | 1.35 ± 0.81 | 2.60 ± 1.17 |
| LINC00861 | 0.21 ± 0.34 | 0.08 ± 0.13 | 0.05 ± 0.07 | 10.34 ± 5.29 | 10.49 ± 3.64 | 9.21 ± 2.00 |
| LINC01550 | 0.03 ± 0.05 | 0.01 ± 0.02 | 0.00 ± 0.00 | 0.02 ± 0.01 | 2.20 ± 0.52 | 1.97 ± 0.21 |
| LINC02295 | 0.00 ± 0.00 | 0.00 ± 0.00 | 0.00 ± 0.00 | 0.00 ± 0.00 | 0.81 ± 0.90 | 0.46 ± 0.28 |
| LINC02361 | 0.72 ± 0.25 | 0.48 ± 0.28 | 0.61 ± 0.43 | 2.30 ± 0.29 | 9.96 ± 3.44 | 5.83 ± 1.50 |
| LINC02273 | 0.03 ± 0.05 | 0.01 ± 0.02 | 0.06 ± 0.02 | 0.27 ± 0.05 | 5.50 ± 0.67 | 3.01 ± 1.35 |
| LINC02446 | 0.15 ± 0.26 | 0.02 ± 0.02 | 0.01 ± 0.01 | 0.40 ± 0.36 | 0.04 ± 0.05 | 15.72 ± 7.31 |

**Table S1.** Myeloid and lymphoid cell specific lincRNAs identified in the current study.

| Gene ID | Name |
|---|---|
| ENSG00000140678 | ITGAX |
| ENSG00000160255 | ITGB2 |
| ENSG00000011600 | TYROBP |
| ENSG00000066336 | SPI1 |
| ENSG00000147872 | PLIN2 |
| ENSG00000111679 | PTPN6 |
| ENSG00000104921 | FCER2 |
| ENSG00000232810 | TNF |
| ENSG00000128604 | IRF5 |
| ENSG00000170909 | OSCAR |
| ENSG00000160294 | MCM3AP |
| ENSG00000185291 | IL3RA |
| ENSG00000143387 | CTSK |
| ENSG00000136826 | KLF4 |
| ENSG00000135363 | LMO2 |
| ENSG00000136869 | TLR4 |
| ENSG00000105383 | CD33 |
| ENSG00000021355 | SERPINB1 |
| ENSG00000105810 | CDK6 |
| ENSG00000150337 | FCGR1A |
| ENSG00000119535 | CSF3R |
| ENSG00000198223 | CSF2RA |
| ENSG00000163220 | S100A9 |
| ENSG00000143546 | S100A8 |
| ENSG00000118513 | MYB |
| ENSG00000090382 | LYZ |

**Table S2.** List of PU.1 target genes.

| Patient Nr. | Gender | Age | Patient group (WHO grade) |
|---|---|---|---|
| 1 | Female | 68 | Control |
| 2 | Female | 46 | Control |
| 3 | Male | 92 | Control |
| 4 | Female | 47 | Control |
| 5 | Male | 74 | Control |
| 6 | Male | 65 | Control |
| 7 | Male | 68 | Control |
| 8 | Male | 83 | COVID-19 (> 4) |
| 9 | Male | 88 | COVID-19 (> 4) |
| 10 | Male | 78 | COVID-19 (3) |
| 11 | Female | 72 | COVID-19 (8) |
| 12 | Male | 70 | COVID-19 (8) |
| 13 | Male | 49 | COVID-19 (8) |
| 14 | Female | 52 | COVID-19 (5) |
| 15 | Male | 55 | COVID-19 (4) |
| 16 | Male | 59 | COVID-19 (4) |
| 17 | Female | 42 | COVID-19 (2) |
| 18 | Female | 71 | COVID-19 (4) |
| 19 | Male | 55 | COVID-19 (4) |

**Table S3.** COVID patient characteristics (cohort 1). Red: patient PBMCs analyzed by single cell RNA-seq.

| Patient Nr. | Gender | Age | Patient group (intensive care unit) |
|---|---|---|---|
| 1 | Female | 86 | Control |
| 2 | Female | 65 | Control |
| 3 | Male | 44 | Control |
| 4 | Male | 42 | Control |
| 5 | Female | 80 | Control |
| 6 | Female | 72 | Control |
| 7 | Female | 26 | Control |
| 8 | Male | 41 | Control |
| 9 | Male | 28 | Control |
| 10 | Male | 70 | Control |
| 11 | Male | 28 | Control |
| 12 | Male | 82 | COVID-19 (ICU) |
| 13 | Female | 58 | COVID-19 |
| 14 | Male | 52 | COVID-19 |
| 15 | Male | 44 | COVID-19 (ICU) |
| 16 | Male | 66 | COVID-19 (ICU) |
| 17 | Male | 23 | COVID-19 (ICU) |
| 18 | Male | 82 | COVID-19 (ICU) |

**Table S4.** COVID patient characteristics (cohort 2). ICU = intensive care unit.

| Patient Nr. | Gender | Age | Patient group (type of infection) |
|---|---|---|---|
| 1 | male | 53 | control |
| 2 | male | 49 | control |
| 3 | male | 38 | control |
| 4 | female | 71 | control |
| 5 | female | 49 | control |
| 6 | male | 62 | control |
| 7 | male | 28 | control |
| 8 | male | 29 | control |
| 9 | female | 27 | control |
| 10 | male | 64 | control |
| 11 | female | 25 | infection (bacterial) |
| 12 | male | 45 | infection (bacterial) |
| 13 | female | 64 | infection (bacterial) |
| 14 | female | 19 | infection (fungal) |
| 15 | female | 64 | infection (fungal) |
| 16 | female | 42 | infection (fungal) |
| 17 | female | 68 | infection (n.d.) |
| 18 | female | 59 | infection (n.d.) |
| 19 | female | 52 | infection (n.d.) |
| 20 | female | 58 | infection (n.d.) |
| 21 | female | 73 | infection (n.d.) |
| 22 | female | 61 | infection (polymicrobial) |

**Table S5.** Characteristics of control and pulmonary infection patients. n.d. = causative pathogen not determined

| Target (purpose) | Oligo name | Oligo sequence |
|---|---|---|
| SparQ MCS (Sanger sequencing) | OBS-0659 | Fwd: AGGAGGATTTGATATTCACCTG |
| | OBS-0660 | Rev: ACCTTCTCTAGGCACCCG |
| pX458 (Sanger sequencing) | OBS-0842 | Fwd: CTGGCCTTTTGCTCACATGT |
| | OBS-0843 | Rev: GTCTGCAGAATTGGCGCAC |
| hPIRAT (qRT-PCR) | OBS-0808 | Fwd: TGAGTGTACTGCCTGGACTCATC |
| | OBS-0809 | Rev: TAAATGAAGCCTGCCACTTCAG |
| hLUCAT1 (qRT-PCR) | OBS-0849 | Fwd: ACCATGTGTCAAGCTCGGATTG |
| | OBS-0850 | Rev: TTGTGGTCTCTGGTGCCAAG |
| AC064805.1 (qRT-PCR) | OBS-0899 | Fwd: TAATAAGCAGCAATTGCAGTTCC |
| | OBS-0900 | Rev: TATCTGCTCCTGAGGCAGAGG |
| LINC02295 (qRT-PCR) | OBS-0935 | Fwd: AAGCTGTGGCTGTTGTCAGC |
| | OBS-0936 | Rev: ACACTTGTCTCAGTAGGCCTGG |
| LINC02446 (qRT-PCR) | OBS-0937 | Fwd: TGTCACCTGTGGACAACTTGC |
| | OBS-0938 | Rev: TTATCTTGACCAGGTGCGAGAC |
| LINC00861 (qRT-PCR) | OBS-0907 | Fwd: AACACTGAGCAATCCTGACCTG |
| | OBS-0908 | Rev: TATCGGTCCTCCACTCTTGTTC |
| hS100A8 (qRT-PCR) | OBS-2077 | Fwd: AGACTGTAGCAACTCTGGCAG |
| | OBS-2078 | Rev: TCCAGCTCGGTCAACATGATG |
| hS100A9 (qRT-PCR) | OBS-2198 | Fwd: ACCAATACTCTGTGAAGCTGG |
| | OBS-2199 | Rev: TCCTCGAAGCTCAGCTGCTTG |
| hCHI3L1 (qRT-PCR) | OBS-2179 | Fwd: AGGGACCCTTGCCTACTATGA |
| | OBS-2180 | Rev: TGGAAGTCATCCAGGTCCAGG |
| hITGAX (qRT-PCR) | OBS-2275 | Fwd: AAGCTGACAGACGTGGTCATC |
| | OBS-2276 | Rev: ATACTGCAGCCTGGAGGAGAG |
| hCXCL2 (qRT-PCR) | OBS-534 | Fwd: CCTGCAGGGAATTCACCTCA |
| | OBS-535 | Rev: CCTTCCTTCTGGTCAGTTGG |
| hIL6 (qRT-PCR) | OBS-15 | Fwd: AATTCGGTACATCCTCGACGG |
| | OBS-16 | Rev: TTGGAAGGTTCAGGTTGTTTTCT |
| hPU.1 (qRT-PCR) | OBS-2399 | Fwd: AGAGCCATAGCGACCATTACTG |
| | OBS-2400 | Rev: ATCTGCTCCAGCTCCATGTG |
| hGAPDH (qRT-PCR) | OBS-0430 | Fwd: CCACATCGCTCAGACACCAT |
| | OBS-0431 | Rev: CGCAACAATATCCACTTTACCAGAG |
| hMALAT1 (qRT-PCR) | OBS-814 | Fwd: AGGTGCTACACAGAAGTGGATTCAG |
| | OBS-815 | Rev: CTTCCCGTACTTCTGTCTTCCAGT |
| hU6 (qRT-PCR) | OBS-0712 | Fwd: GCTTCGGCAGCACATATACTAAAAT |

| | | |
|---|---|---|
| | OBS-0713 | Rev: ATATGGAACGCTTCACGAATTTG |
| hS100A8 promoter (qRT-PCR) | OBS-3211 | Fwd: AGCATCCACTTCCTATTCCTGC |
| | OBS-3212 | Rev: TAAGGATTTGGGTAGCATGGAGG |
| hS100A9 promoter (qRT-PCR) | OBS-3213 | Fwd: TGAACTAAACAACCAGCTTCCTCC |
| | OBS-3214 | Rev: TGAGCAGTGTGGTAATGCTGC |
| hREXOL1P peak1 (qRT-PCR) | OBS-3215 | Fwd: ACAGTGAGTTGGTCAAATGCTCC |
| | OBS-3216 | Rev: ACAGCATAGGTTGAGAAGCTGTTAC |
| hREXOL1P peak2 (qRT-PCR) | OBS-3221 | Fwd: TTCTCCACACTGTCAGGAGC |
| | OBS-3222 | Rev: AGGTGTAGGAAGCCATACACTG |
| hREXOL1P peak3 (qRT-PCR) | OBS-3202 | Fwd: CCACAGCCAATATCATACTGAATGG |
| | OBS-3203 | Rev: ACAAACAGGGACAATTTGACTTCCTC |
| hREXOL1P peak4 (qRT-PCR) | OBS-3204 | Fwd: GCTGAAGTTGCTTATCAGCTTAAGG |
| | OBS-3205 | Rev: ACATAGTGTTGGAACTTCTGGCC |
| hPIRAT (genomic PCR) | OBS-1895 | Fwd: GCATCTGCATGGCAGAGTTC |
| | OBS-1239 | Rev: TATGGCTCTTGCAATTAATCCTG |
| PIRAT 3’ RACE | OBS-1058 | AGAGTGAGTGTACTGCCTGGACTCATCAC |
| PIRAT 5’ RACE | OBS-1059 | ACATAAAGATGTGGACAGTCACCATTGTC |
| PU.1 gRNA inserts | OBS-2281 | Sense strand: CACCGAAATCTCTTGCGCTACATAC |
| | OBS-2282 | Antisense strand: AAACGTATGTAGCGCAAGAGATTTC |

**Table S6.** PCR and sequencing oligonucleotides used in the present study.

| Target (purpose) | Oligo name | Oligo sequence |
|---|---|---|
| PIRAT | OBS-2890 | TCCAATTAGCCAAGAGCATG |
| PIRAT | OBS-2891 | CTTTCACTGTTGTGACCATT |
| PIRAT | OBS-2892 | GTTTTCAGTCTAAAGCCAGT |
| PIRAT | OBS-2893 | CTCATCTCTTCTTTCTGTTA |
| PIRAT | OBS-2894 | ATGCCAAAGGCCAGACAAAC |
| PIRAT | OBS-2895 | AAAGTGATGAGTCCAGGCAG |
| PIRAT | OBS-2896 | CTGCCACTTCAGTGTAGAAA |
| PIRAT | OBS-2897 | TGTCCAAAACGTGGGTGTTA |
| PIRAT | OBS-2898 | ATTACCTCAGATCACGGTTG |
| None (control ChIRP) | OBS-2899 | CACTATGGAAAGGCGGCTTC |
| None (control ChIRP) | OBS-2900 | GATTTCGGTCTGTACGGCTA |
| None (control ChIRP) | OBS-2901 | TTACATGGTCCTAATCGGCT |
| None (control ChIRP) | OBS-2902 | GCTGTTACCTTCCACGCCGG |
| None (control ChIRP) | OBS-2903 | ATACGATCGGACAGCCTTGT |
| None (control ChIRP) | OBS-2904 | TGCACAATTGATGTTCCGAT |
| None (control ChIRP) | OBS-2905 | GACGCCTAGACGTATACTAG |
| None (control ChIRP) | OBS-2906 | GTGTGTGCTATTAGAAGCGG |
| None (control ChIRP) | OBS-2907 | AAGCGACCCTGACAGTGCGA |
| None (control ChIRP) | OBS-2908 | AGCAAACACGTCGAGCAAAT |

**Table S7.** ChIRP oligonucleotides used in the present study.

| Specificity | Source | Class | Conjugate | Supplier | Catalog nr. | Application |
|---|---|---|---|---|---|---|
| PU.1 (A7) | mouse | IgG1 | - | Santa Cruz | sc-365208 | Western Blot, IP |
| PU.1 (C3) | mouse | IgG | - | Santa Cruz | sc-390405 | IP |
| Flag | mouse | IgM2 | - | Sigma-Aldrich | F1804 | IP |
| Anti-mouse | goat | IgG | HRP | Santa Cruz | sc-2005 | Western Blot |
| CD14 | mouse | IgG1 | FITC | eBioscience | 11-0149-42 | FACS |
| CD66b | mouse | IgM | APC | eBioscience | 17-0666-42 | FACS |
| CD4 | mouse | IgG2b | PE | eBioscience | 12-0048-41 | FACS |
| CD8 | mouse | IgG1 | FITC | eBioscience | 11-0087-42 | FACS |
| CD19 | mouse | IgG1 | FITC | eBioscience | 11-0199-41 | FACS |
| CD56 | mouse | IgG | APC | eBioscience | 17-0567-41 | FACS |
| CD11c | mouse | IgG1 | APC | eBioscience | 17-0116-42 | FACS |

**Table S8.** Antibodies used in the present study.
